# Localized mRNA translation mediates maturation of cytoplasmic cilia in *Drosophila* spermatogenesis

**DOI:** 10.1101/2020.04.21.054247

**Authors:** Jaclyn M Fingerhut, Yukiko M Yamashita

**Affiliations:** Cellular and Molecular Biology Program, University of Michigan, Ann Arbor, MI 48109, USA; Life Sciences Institute, University of Michigan, Ann Arbor, MI 48109, USA; Department of Cell and Developmental Biology, University of Michigan, Ann Arbor, MI 48109, USA; Howard Hughes Medical Institute, University of Michigan, Ann Arbor, MI 48109, USA

## Abstract

Cytoplasmic cilia, a specialized type of cilia in which the axoneme resides within the cytoplasm rather than within the ciliary compartment, are proposed to allow the efficient assembly of very long cilia. Despite being found diversely in male gametes (e.g. *Plasmodium* microgametocytes and human and *Drosophila* sperm), very little is known about cytoplasmic cilia assembly. Here we show that a novel RNP granule containing the mRNAs for axonemal dynein motor proteins becomes highly polarized to the distal end of the cilia during cytoplasmic ciliogenesis in *Drosophila* sperm. This allows for the localized translation of these axonemal dyneins and their incorporation into the axoneme directly from the cytoplasm. We found that this RNP granule contains the proteins Reptin and Pontin, loss of which perturbs granule formation and prevents incorporation of the axonemal dyneins, leading to sterility. We propose that cytoplasmic cilia require the local translation of key protein constituents such that these proteins are incorporated efficiently into the axoneme.

**Author Summary:** Cytoplasmic cilia, which are found in human and *Drosophila* sperm, are unique in that the axoneme is exposed to the cytoplasm. The authors show that a novel RNP granule containing axonemal dynein mRNAs facilitates localized translation of these axonemal proteins, facilitating cytoplasmic cilia formation.

## Introduction

Cilia are microtubule-based structures present on the surface of many cells. These specialized cellular compartments can be non-motile primary cilia that largely function in signaling, or motile cilia that can either move extracellular materials (e.g. lung multiciliated cells) or allow for cell motility (e.g. *Chlamydomonas* flagellum, sperm in many species) (Ishikawa and Marshall, 2011). It is well established that most cilia are separated from the bulk cytoplasm (Fig. 1 A), which serves to concentrate signaling molecules for rapid response to extracellular signals received by the cilia, and that the ciliary gate at the base of the cilia forms a diffusion barrier, through which molecules must be selectively transported (Reiter et al., 2012; Wheway et al., 2018). However, recent studies identified an additional type of cilia, called cytoplasmic cilia, in which the axoneme (the microtubule-based core of the cilia) is exposed to the cytoplasm (Fig. 1 A) (Avidor-Reiss et al., 2017; Avidor-Reiss and Leroux, 2015; Dawson and House, 2010; Fawcett et al., 1970; Sinden et al., 1976; Tates, 1971; Tokuyasu, 1975). Cytoplasmic cilia are found in human and *Drosophila* sperm as well as in *Plasmodium* and *Giardia*. There are two proposed advantages to cytoplasmic cilia: 1) faster assembly as the cell does not need to rely on ciliary transport mechanisms, allowing for the assembly of longer cilia, and 2) proximity to mitochondria for energy (Avidor-Reiss and Leroux, 2015; Sinden et al., 2010). Despite being found across diverse taxa, very little is known about how cytoplasmic cilia are assembled and whether their assembly bears similarity to that of traditional compartmentalized cilia (Desai et al., 2018).

**Figure 1:**
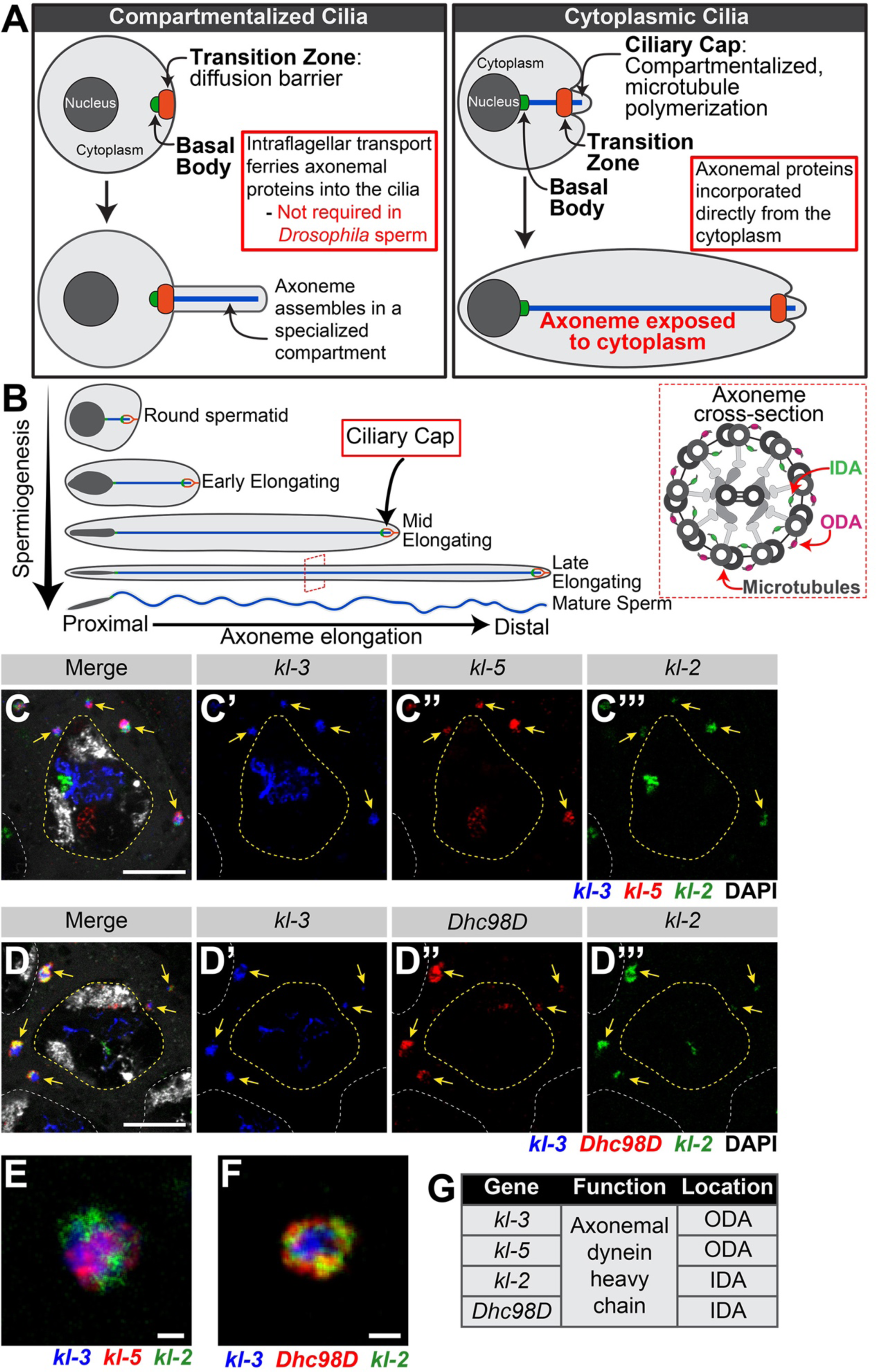
Axonemal dynein heavy chain mRNAs colocalize in an RNP granule in spermatocytes. **(A)** Diagram comparing and contrasting traditional compartmentalized cilia and cytoplasmic cilia. Nucleus (dark gray), cytoplasm (light gray), basal body (green), transition zone (orange) and axoneme (blue). **(B)** Diagram of *Drosophila* spermiogenesis with stages of spermatid elongation. Nucleus (dark gray), cytoplasm (light gray), basal body & ring centriole (green), ciliary cap (orange) and axoneme (blue). Axoneme cross section image showing location of axonemal dynein arms. Microtubules and other structural components (gray), inner dynein arm (green) and outer dynein arm (magenta). **(C and D)** smFISH against axonemal dynein heavy chain transcripts in SCs showing *kl-3, kl-5*, and *kl-2* mRNAs (C) or *kl-3, kl-2*, and *Dhc98D* mRNAs (D) in kl-granules. *kl-3* (blue), *kl-2* (green), *kl-5* (red, C), *Dhc98D* (red, D), DAPI (white), kl-granules (yellow arrows), SC nuclei (yellow dashed line), neighboring SC nuclei (white dashed line). Bar: 10μm. **(E and F)** smFISH against *kl-3, kl-5*, and *kl-2* (E) or *kl-3, kl-2*, and *Dhc98D* (F) showing a single kl-granule. *kl-3* (blue), *kl-2* (green), *kl-5* (red, E), *Dhc98D* (red, F). Bar: 1μm. **(G)** Table listing the genes focused on in this study, their function and their localization within the axoneme.

Cytoplasmic ciliogenesis has been proposed to occur in two stages (Fig. 1 A) (Avidor-Reiss and Leroux, 2015). In the first stage, microtubules are polymerized in a small compartmentalized region, which is similar to canonical compartmentalized cilia, at the most distal end of the cilia (Gottardo et al., 2013; Tokuyasu, 1975). This region is gated by a transition zone (Caudron and Barral, 2009; Kwitny et al., 2010; Vieillard et al., 2016). This entire compartmentalized region, called the ciliary cap or the growing end, migrates away from the basal body, which is docked at the nuclear membrane (Basiri et al., 2014; Fawcett et al., 1970). The ciliary cap does not change in size as the cilia elongates. The continued polymerization of microtubules inside the ciliary cap displaces recently synthesized microtubules out of the compartmentalized region, exposing them to the cytoplasm (Fig. 1 A). The second stage is axoneme maturation, in which additional axonemal proteins (e.g. axonemal dyneins, the motor proteins that confer motility by allowing axonemal microtubules to slide against each other, Fig. 1 B) are added to the bare microtubule structure after it emerges from the ciliary cap (Tates, 1971; Tokuyasu, 1975). Axoneme maturation was inferred to occur in the cytoplasm based on the dispensability of ciliary transport mechanisms and the inefficiency of relaying on diffusion through the transition zone (Avidor-Reiss and Leroux, 2015; Breslow et al., 2013; Briggs et al., 2004; Han et al., 2003; Hoeng et al., 2008; Kee et al., 2012; Lin et al., 2013; Sarpal et al., 2003). However, how this maturation process occurs in the cytoplasmic compartment to allow for cytoplasmic cilia formation remains unknown.

*Drosophila* spermatogenesis provides an excellent model for the study of cytoplasmic ciliogenesis (Fig. 1 B), owing to rich cytological knowledge of spermatogenesis and the conservation of almost all known ciliary proteins (Zur Lage et al., 2019). Developing spermatids elongate from 15μm to 1,900μm (1.9mm) (Tates, 1971; Tokuyasu, 1975). Within mature sperm, the cytoplasmic cilia are 1,800μm and the ciliary caps (the compartmentalized region) are only ∼2μm. Ciliogenesis starts in premeiotic spermatocytes (SCs), which assemble short primary (compartmentalized) cilia (Fabian and Brill, 2012; Gottardo et al., 2013; Riparbelli et al., 2012; Tates, 1971). Prior to axoneme elongation, these primary cilia, which were docked at the plasma membrane in SCs, invaginate, forming the ciliary cap. During axoneme elongation, the majority of the length of the cilia will be exposed to the cytoplasm, as described above. Accordingly, axoneme assembly in *Drosophila* does not require intraflagellar transport (IFT) (Han et al., 2003; Sarpal et al., 2003), the process used by traditional compartmentalized cilia to ferry axonemal proteins from the cytoplasm into the ciliary compartment for incorporation (Rosenbaum and Witman, 2002). Other cytoplasmic cilia have been found not to require IFT for their assembly (Avidor-Reiss and Leroux, 2015; Briggs et al., 2004; Hoeng et al., 2008), leading to the appreciation of a distinct type of cilia: based on the dispensability of IFT, it was postulated that axoneme maturation must occur in the cytoplasm, hence the term ‘cytoplasmic cilia’.

It has long been known that SCs transcribe almost all genes whose protein products are needed post-meiotically and that these mRNAs may not be translated until days later when proteins are needed (Barckmann et al., 2013; Olivieri and Olivieri, 1965). We previously showed that the Y-linked testis-specific axonemal dynein heavy chain genes *kl-3* and *kl-5*, as well as the testis-specific axonemal dynein intermediate chain *Dic61B*, are transcribed in SCs (Fingerhut et al., 2019). However, axoneme elongation does not begin until after meiosis, suggesting that these mRNAs may not be translated until later in development. Intriguingly, we previously showed that *kl-3* and *kl-5* mRNAs localize to cytoplasmic granules in SCs. Ribonucleoprotein (RNP) granules (e.g. stress granules, P granules and germ granules) are known to play critical roles in mRNA regulation, such as mediating the subcellular localization of mRNAs and controlling the timing of translation (Anderson and Kedersha, 2009; Buchan, 2014; Medioni et al., 2012). Therefore, we decided to investigate the role of this novel RNA granule in the translational regulatory mechanisms that ensure proper axoneme assembly and found that it plays an essential role in the incorporation of axonemal proteins, providing the first insights into the molecular mechanism of cytoplasmic cilia maturation. We show four axonemal dynein heavy chain mRNAs, including *kl-3* and *kl-5*, colocalize in these novel granules in late SCs along with the AAA+ (ATPases Associated with diverse cellular Activities) proteins Reptin (Rept) and Pontin (Pont). These RNP granules are segregated during the meiotic divisions and localize to the distal end of the cytoplasmic compartment as the axoneme elongates during spermiogenesis. We further show that Rept and Pont are required for RNP granule formation, and that RNP granule formation is necessary for robust translation and incorporation of the axonemal dynein proteins into the axoneme. We propose that cytoplasmic cilia maturation relies on the local translation of axonemal components such that they can be incorporated into the bare microtubule structure as it emerges from the ciliary cap.

## Results

### Axonemal dynein heavy chain mRNAs colocalize in RNP granules in spermatocytes

In our previous study, we analyzed the expression of the Y-linked axonemal dynein genes *kl-3* and *kl-5* and showed that these two mRNAs localized to cytoplasmic granules in late SCs (Fingerhut et al., 2019). Using single molecule RNA fluorescent *in situ* hybridization (smFISH), we found that mRNAs for four testis-specific axonemal dynein heavy chain genes (the Y-chromosome genes *kl-2, kl-3*, and *kl-5*, as well as the autosomal gene *Dhc98D* (Carvalho et al., 2000; Goldstein et al., 1982; Hardy et al., 1981; Zur Lage et al., 2019)) colocalize together within RNP granules in the cytoplasm of late SCs, with each SC containing several of these cytoplasmic granules (Fig. 1 C and D). We termed these granules “kl-granules” after the three Y-linked constituent mRNAs. It should be noted that robust transcription of these genes is still ongoing in SC nuclei (visible as bright nuclear signal, Fig. 1 C and D) but these are nascent transcripts that still contain intronic RNA, whereas the kl-granules in the cytoplasm do not contain intronic RNA, as we showed previously (Fingerhut et al., 2019). The present study focuses the fate of these cytoplasmic RNPs that contain mature mRNA. mRNAs within a kl-granule are spatially sub-organized: *kl-3* and *kl-5* mRNAs, which encode outer dynein arm (ODA) dynein heavy chain proteins, cluster together in the core of the kl-granule while *kl-2* and *Dhc98D* mRNAs, which encode inner dynein arm (IDA) dynein heavy chain proteins, localize peripherally (Fig. 1 E – G). This is similar to the sub-compartmentalization observed in other RNP granules, including the germ granules in the *Drosophila* ovary, stress granules, P granules and nucleoli (Boisvert et al., 2007; Jain et al., 2016; Trcek et al., 2015; Wang et al., 2014). We noted that kl-granule formation is unlikely to be dependent upon any one mRNA constituent as RNAi mediated knockdown of *kl-3, kl-5, kl-2*, or *Dhc98D* (*bam-gal4>UAS-kl-3*^*TRiP.HMC03546*^ or *bam-gal4>UAS-kl-5*^*TRiP.HMC03747*^ or *bam-gal4>UAS-kl-2*^*GC8807*^ or *bam-gal4>UAS-Dhc98D*^*TRiP.HMC06494*^) did not perturb granule formation despite efficient knockdown (Fig. S1).

We conclude that mRNAs for the testis-specific axonemal dynein heavy chains localize to novel RNP granules, which we termed kl-granules, in late SCs.

### The kl-granules segregate during the meiotic divisions and localize to the distal end of elongating spermatids

As the kl-granules contain mRNAs for axonemal proteins that are only necessary for spermiogenesis, we next followed the fate of the kl-granules through meiosis and into spermiogenesis. The kl-granules segregate through the two sequential meiotic divisions (Fig. 2 A) such that each resulting haploid spermatid receives a relatively equal amount of kl-granule (Fig. 2 B). Upon completion of meiosis, the resultant spermatids are interconnected due to incomplete cytokinesis during the four mitotic divisions that occur early in germ cell development and the two meiotic divisions, forming a cyst of 64 spermatids (Fuller, 1993; Hime et al., 1996). As the axoneme starts to elongate within each spermatid, the nuclei cluster to the proximal end of the cyst while the axoneme elongates unidirectionally away from the nuclei with the ciliary caps clustered at the distal end of the cyst (Fig. 1 B) (Fabian and Brill, 2012). Strikingly, we found that the kl-granules become localized to the distal end of elongating spermatid cysts (Fig. 2 C). This polarized localization remains as the axoneme continues to elongate (Fig. 2 D and E). At later stages of elongation, the kl-granules begin to dissociate and the mRNAs become more diffusely localized at the distal end (Fig. 2 D and E). Interestingly, some mRNAs dissociate from the kl-granules before others: *kl-3* and *kl-5* mRNAs (encoding ODA proteins) dissociate earlier than *kl-2* and *Dhc98D* mRNAs (encoding IDA proteins) (Fig. 2 D and F). It is of note that the differential timing of dissociation correlates with the sub-compartmentalization of constituent mRNAs described above: *kl-3* and *kl-5* localize to the core of the kl-granules and dissociate first (Fig. 1 E), whereas *kl-2* and *Dhc98D* localize to the periphery of the kl-granules (Fig. 1 F) and dissociate later. These results show that kl-granules exhibit stereotypical localization to the growing end of spermatids after being segregated during meiosis, implying that programmed positioning of the kl-granules may play an important role during spermatid elongation and axoneme maturation.

**Figure 2:**
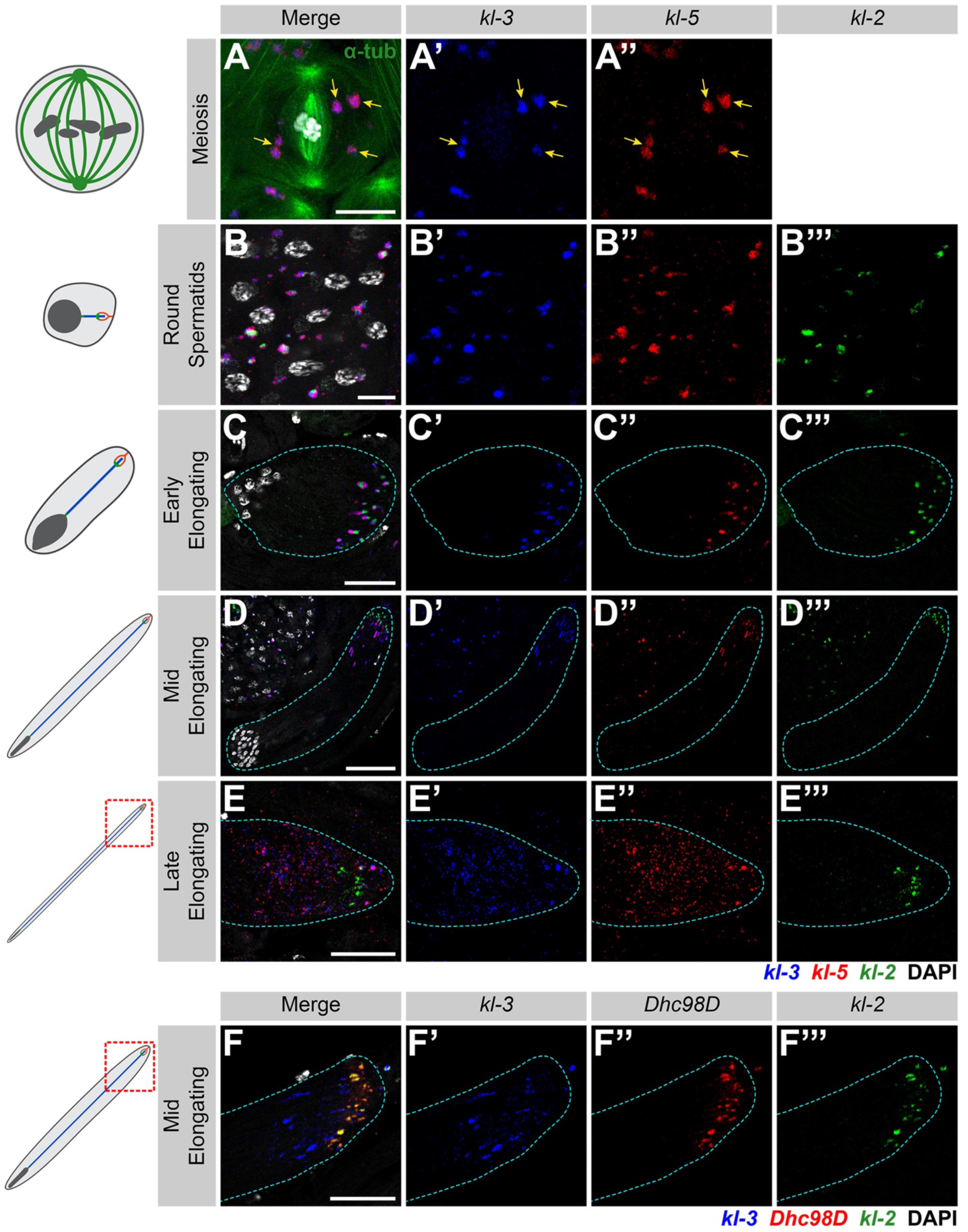
kl-granules segregate during the meiotic divisions and localize to the distal end of elongating spermatids. **(A)** smFISH against *kl-3* and *kl-5* during meiosis. *kl-3* (blue), *kl-5* (red), α-tubulin-GFP (green), DAPI (white) and kl-granules (yellow arrows). Bar: 10μm. **(B – E)** smFISH against *kl-3, kl-5*, and *kl-2* during spermiogenesis. The round spermatid (B), early elongating spermatid (C), mid elongating spermatid (D) and late elongating spermatid (E) stages are shown. *kl-3* (blue), *kl-5* (red), *kl-2* (green), DAPI (white), spermatid cyst (cyan dashed line). Bar: 10μm (B), 25μm (C and E) or 50μm (D). **(F)** smFISH against *kl-3, Dhc98D*, and *kl-2* in mid elongating spermatids. *kl-3* (blue), *Dhc98D* (red), *kl-2* (green), DAPI (white), spermatid cyst (cyan dashed line). Bar: 25μm.

### The AAA+ proteins Reptin and Pontin colocalize with the kl-granules

To further understand how kl-granules form and their potential function, we sought to identify a protein(s) that localize to the kl-granules. In our previous study, we screened for proteins involved in the expression of the Y-linked axonemal dynein genes (Fingerhut et al., 2019). Reptin (Rept) and Pontin (Pont), two AAA+ proteins (Puchades et al., 2020), were included in this screen because of their high expression in the testis and their involvement in RNP complex formation in other systems (Mao and Houry, 2017; Robinson et al., 2013). Also, studies in *Drosophila*, mouse, zebrafish, *Chlamydomonas* and Xenopus have specifically implicated Rept and Pont in axoneme/motile cilia assembly and/or sperm motility, although the underlying mechanism remains unknown (Dafinger et al., 2018; Huizar et al., 2018; Li et al., 2017; Stolc et al., 2005; Tammana and Tammana, 2017; Zhao et al., 2013; Zur Lage et al., 2018).

We found that Rept and Pont colocalize in cytoplasmic granules in SCs through elongating spermatids (Fig. 3 A and B). Immunofluorescent staining combined with smFISH (IF-smFISH) showed that Rept and Pont colocalize with the kl-granules. Pont first colocalizes with *Dhc98D* mRNA in early SCs (Fig. 3 C) and with all other kl-granule constituent mRNAs in later SCs (Fig. 3 D) and throughout spermatid elongation (Fig. 3 E). Close examination of the kl-granules in late SCs revealed that Pont is not evenly distributed within a kl-granule and rather concentrates near the core with *kl-3* and *kl-5* mRNAs (Fig. 1 E and Fig. 3 F). In contrast, *kl-2* and *Dhc98D* mRNAs occupy the periphery of the kl-granule (Fig. 1 F), where Pont is less concentrated.

**Figure 3:**
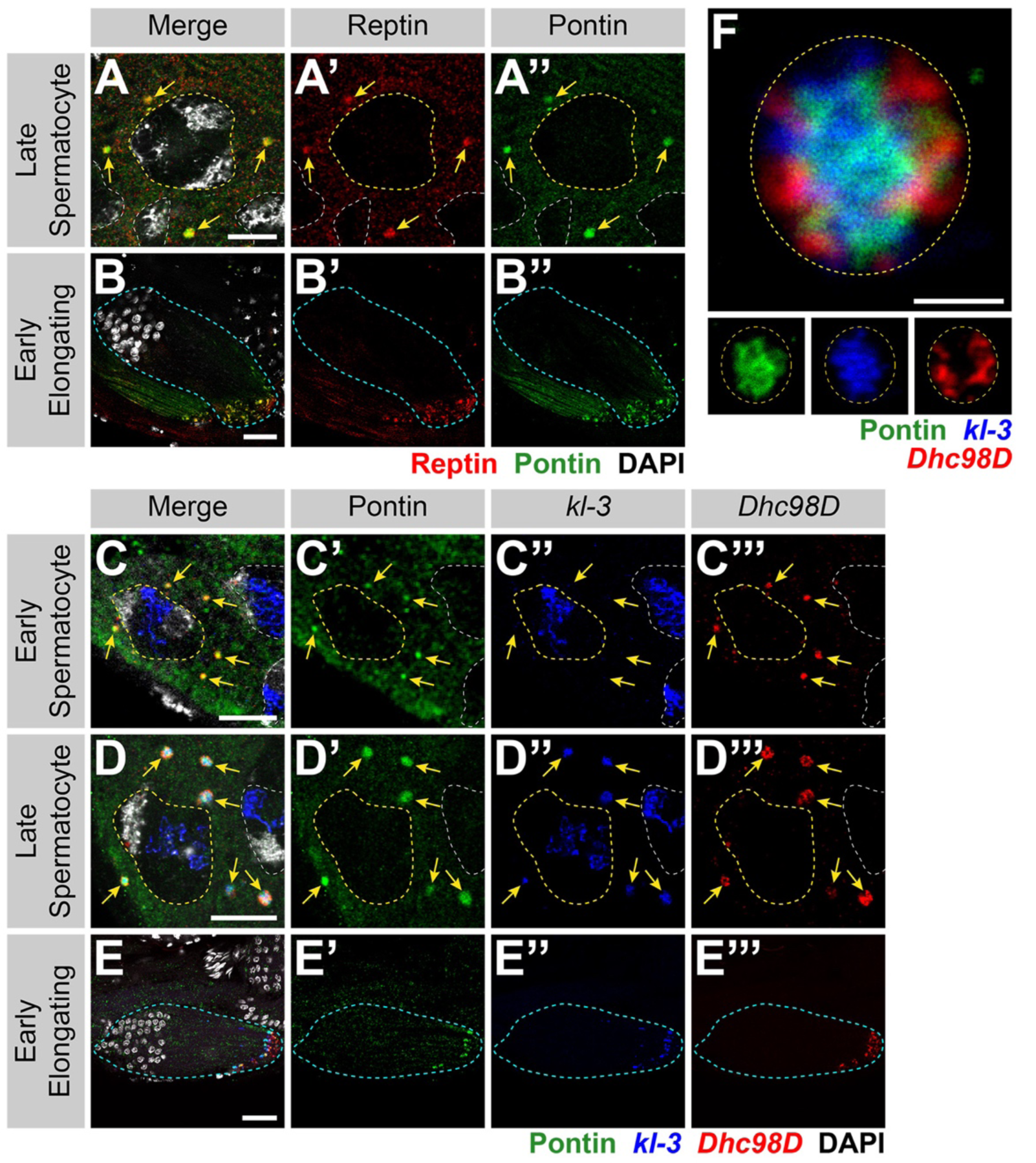
Reptin and Pontin colocalize with the kl-granules. **(A and B)** Rept and Pont colocalization in SCs (A) and early elongating spermatids (B). Rept (red), Pont (green), DAPI (white), SC nuclei (yellow dashed line, A), neighboring SC nuclei (white dashed line, A), kl-granules (yellow arrows, C) spermatid cyst (cyan dashed line, B). Bar: 10μm (A) or 25μm (B). **(C – E)** IF-smFISH for Pont protein and *kl-3* and *Dhc98D* mRNAs in early SCs (C), late SCs (D) and early elongating spermatids (E). Pont (green), *kl-3* (blue), *Dhc98D* (red), DAPI (white), SC nuclei (yellow dashed line, C and D), neighboring SC nuclei (white dashed line, C and D), kl-granules (yellow arrows, C and D) spermatid cyst (cyan dashed line, E). Bar: 10μm (C and D) or 25μm (E). **(F)** IF-smFISH for Pont protein and *kl-3* and *Dhc98D* mRNAs in a single kl-granule. Pont (green), *kl-3* (blue), *Dhc98D* (red) and kl-granule boundary (yellow dashed line). Bar: 1μm.

We conclude that Rept and Pont localize to the kl-granules together with axonemal dynein heavy chain mRNAs. It is interesting to note that previous studies have proposed that Rept and Pont function as chaperones in the assembly of axonemal dynein motors (complexes containing a combination of dynein heavy, intermediate, and light chains) (Huizar et al., 2018; Li et al., 2017; Zur Lage et al., 2018). It remains unknown whether previously reported Rept- and Pont-containing chaperon complexes also contain mRNA (see Discussion).

### Reptin and Pontin are required for kl-granule assembly

To explore the function of Rept and Pont in kl-granule formation, we performed RNAi mediated knockdown of either *rept* or *pont* (*bam-gal4>UAS-rept*^*KK105732*^ or *bam-gal4>UAS-pont*^*KK101103*^). In addition to eliminating the targeted protein, depletion of *rept* resulted in loss of Pont and vice versa, reminiscent of findings from previous studies, likely because these proteins stabilize each other as components of the same complex (Fig. S2) (Gorynia et al., 2011; Li et al., 2017; Rivera-Calzada et al., 2017; Venteicher et al., 2008).

We next determined whether Rept and Pont are needed for kl-granule assembly. Indeed, knockdown of *rept* or *pont* resulted in disruption of the kl-granules. smFISH clearly detected the presence of dispersed *kl-3* and *kl-5* mRNAs in late SCs, suggesting that *rept* and *pont* are required for kl-granule formation but not for the stability of the constituent mRNAs (Fig. 4 A – C, note that nuclear signal was oversaturated in order to focus on the dispersed cytoplasmic signal). This effect was more pronounced in elongating spermatids where *kl-3* and *kl-5* mRNAs were diffuse throughout the entire cyst in the RNAi conditions (Fig. 4 D – F). RT-qPCR confirmed that mRNA levels were not reduced compared to cross-sibling controls (Fig. 4 G and H), demonstrating that kl-granule formation is not required for mRNA stability. This is in accordance with observations in other systems that suggest that RNA granule formation is not required for mRNA stability and may be more important for mRNA localization or translation (Bley et al., 2015; Lee et al., 2020).

**Figure 4:**
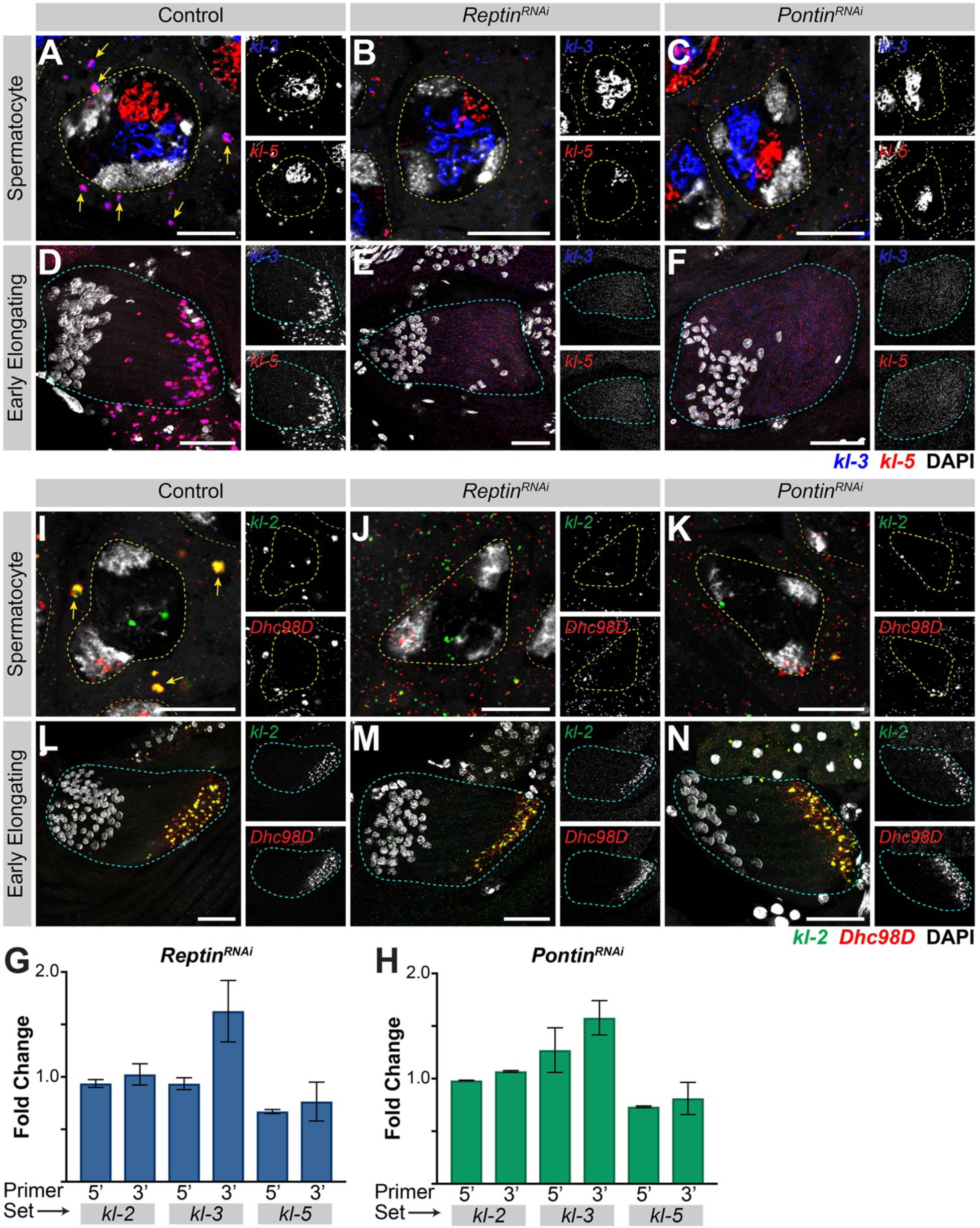
Reptin and Pontin are required for kl-granule assembly. **(A – F)** smFISH against *kl-3* and *kl-5* in control (A and D), *rept* RNAi (*bam-gal4>UAS-rept*^*KK105732*^, B and E) or *pont* RNAi (*bam-gal4>UAS-pont*^*KK101103*^, C and F) SCs (A – C, single z plane) and early elongating spermatids (D – F, z-projection). *kl-3* (blue), *kl-5* (red), DAPI (white), SC nuclei (yellow dashed lines), neighboring SC nuclei (narrow yellow dashed lines), SC kl-granules (yellow arrows) and spermatid cyst (cyan dashed line). Bar: Bar: 10μm (A – C) or 25μm (D – F). **(G and H)** RT-qPCR in *rept* RNAi (*bam-gal4>UAS-rept*^*KK105732*^, G) or *pont* RNAi (*bam-gal4>UAS-pont*^*KK101103*^, H) for *kl-3, kl-5*, and *kl-2* using the indicated primer sets (see Table S1). Data was normalized to GAPDH and sibling controls. **(I – N)** smFISH against *kl-2* and *Dhc98D* in control (I and L), *rept* RNAi (*bam-gal4>UAS-rept*^*KK105732*^, J and M) or *pont* RNAi (*bam-gal4>UAS-pont*^*KK101103*^, K and N) SCs (I – K, single z plane) and early elongating spermatids (L – N, z-projection). *kl-2* (green), *Dhc98D* (red), DAPI (white), SC nuclei (yellow dashed lines), neighboring SC nuclei (narrow yellow dashed lines), SC kl-granules (yellow arrows) and spermatid cyst (cyan dashed line). Bar: Bar: 10μm (I – K) or 25μm (L – N).

Interestingly, knockdown of *rept* or *pont* had a somewhat different effect on *kl-2* and *Dhc98D* mRNAs. smFISH for *kl-2* and *Dhc98D* following RNAi of either *rept* or *pont* showed loss of kl-granule localization in late SCs similar to that seen for *kl-3* and *kl-5* (Fig. 4 I – K). However, in elongating spermatids, *kl-2* and *Dhc98D* mRNAs appeared to localize properly at the distal end of the cyst (Fig. 4 L – N). Considering that Pont primarily colocalized with *kl-3* and *kl-5* mRNAs (Fig. 3 F), this may suggest that other proteins participate in localizing *kl-2* and *Dhc98D* mRNAs to the kl-granule.

In conclusion, Rept and Pont are critical for assembling the kl-granules.

### kl-granule assembly is required for efficient Kl-3 translation and sperm motility

Previous studies in *Drosophila* and mouse demonstrated that Rept and Pont are required for male fertility (Li et al., 2017; Zur Lage et al., 2018). We confirmed that seminal vesicles, where mature motile sperm are stored after exiting the testis, were empty in *rept* or *pont* RNAi testes (Fig. 5 A – C), as was observed for *kl-3, kl-5, kl-2*, or *Dhc98D* RNAi testes (Fig. S3) (Fingerhut et al., 2019; Zur Lage et al., 2018).

**Figure 5:**
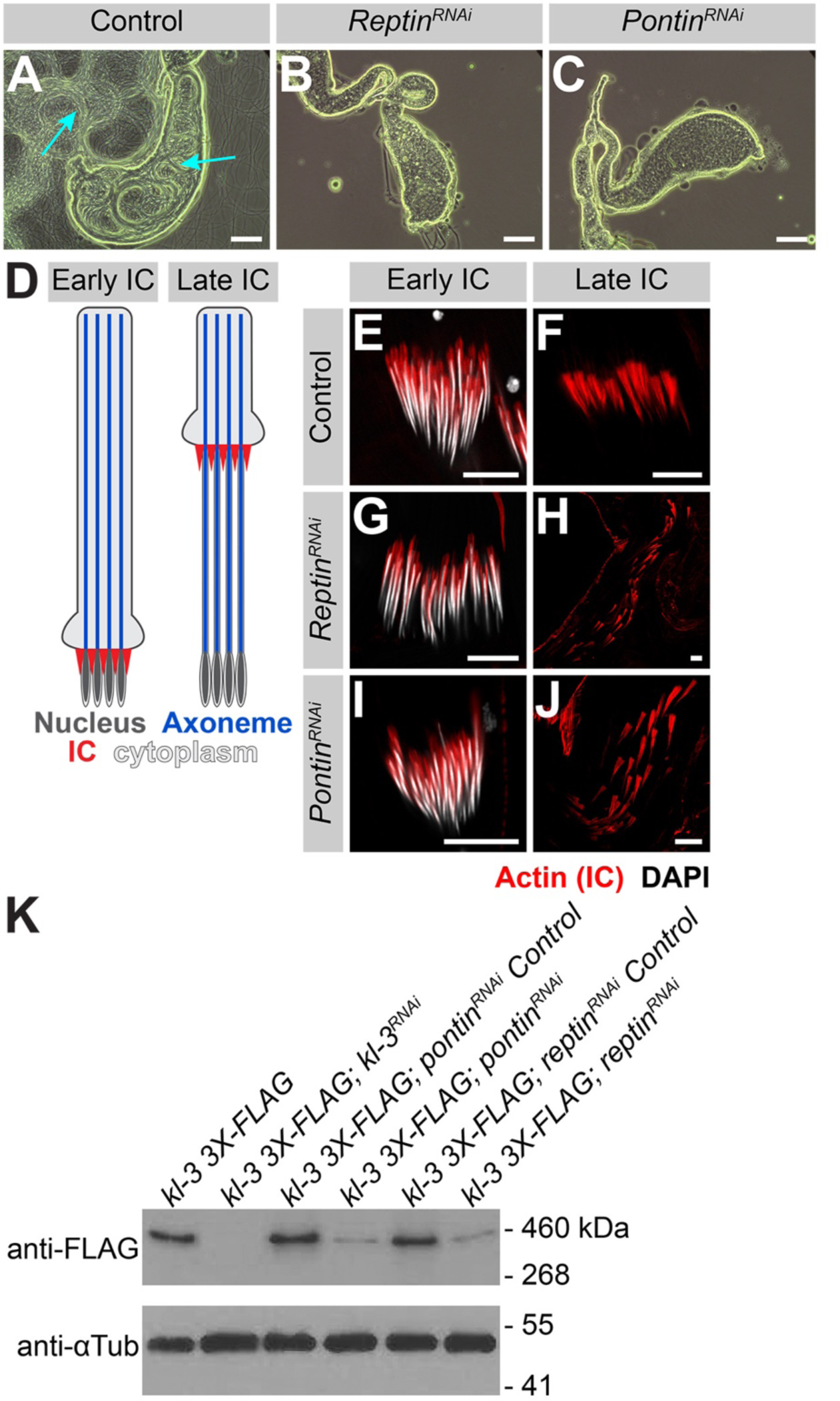
kl-granule assembly is required for efficient Kl-3 translation and sperm motility. **(A – C)** Phase contrast images of seminal vesicles in control (A), *rept* RNAi (*bam-gal4>UAS-rept*^*KK105732*^, B) and *pont* RNAi (*bam-gal4>UAS-pont*^*KK101103*^, C). Mature sperm (cyan arrows). Bar: 100μm. **(D)** Schematic of IC progression during individualization. Nucleus (dark gray), axoneme (blue), ICs (red) and cytoplasm (light gray). **(E – J)** Phalloidin staining of early and late ICs in the indicated genotypes. Phalloidin (Actin, red) and DAPI (white). Bar 10μm. **(K)** Western blot for Kl-3-3X FLAG in the indicated genotypes.

We further characterized the sterility phenotype of *rept* and *pont* RNAi testes and found that spermiogenesis fails during individualization. As sperm develop as cysts, the process of individualization removes excess cytoplasm from the spermatids and separates the cyst into individual sperm via actin-rich individualization complexes (ICs) (Fabian and Brill, 2012). The ICs form around the nuclei at the proximal end of the cyst and progress evenly towards the distal end (Fig. 5 D). It is well established that defects in axoneme assembly, including loss of axonemal dynein motor proteins, perturb IC progression (Fatima, 2011; Fingerhut et al., 2019; Wang et al., 2019). We found that RNAi-mediated knockdown of *rept* or *pont*, does not affect IC assembly but does result in disorganized IC progression (Fig. 5 E – J), as is observed following knockdown of *kl-3, kl-5, kl-2*, or *Dhc98D* (Fig. S3) (Fingerhut et al., 2019).

As previous studies have implicated Rept and Pont in male fertility and axonemal dynein motor assembly, and the observed individualization defects are characteristic of axonemal defects, we analyzed Kl-3 protein levels following *rept* or *pont* RNAi. Western blotting using total testis extracts revealed that Kl-3 protein levels are drastically reduced following knockdown of *rept* or *pont* (Fig. 5 K). Taken together, our results demonstrate that Rept and Pont are required for mRNAs to congress in the kl-granule, which in turn is required for efficient translation. This defect in axonemal dynein expression is the likely cause of sterility in *rept* and *pont* RNAi testes.

### kl-granule formation and localization are required for cytoplasmic cilia maturation

Precise mRNA localization and localized translation are widely utilized mechanisms to ensure that proteins are concentrated where they are needed (Glock et al., 2017; Medioni et al., 2012). As described above, the kl-granules localize to the distal end of elongating spermatids (Fig. 2) where bare axonemal microtubules are first exposed to the cytoplasm after being displaced from the ciliary cap as new microtubules are polymerized. We therefore postulated that the kl-granule may function in cytoplasmic cilia maturation. We first determined whether the kl-granules localize within the ciliary cap or within the cytoplasmic compartment. By using Unc-GFP to mark the ring centriole, a structure at the base of the ciliary cap at the boundary between the cytoplasmic and compartmentalized regions (Baker et al., 2004; Phillips, 1970), we found that the kl-granules are located within the cytoplasmic compartment, immediately proximal to the ciliary cap (Fig. 6 A), suggesting that this may be the site of localized Kl-3 translation. Indeed, we not only found that FLAG-tagged Kl-3 protein (expressed from the endogenous locus, see Methods) occupies the same region proximal to the ciliary cap as the kl-granules but that Kl-3 protein is restricted to the cytoplasmic compartment while the microtubules extend into the compartmentalized compartment (i.e. the ciliary cap) (Fig. 6 B). These results indicate that while the axonemal microtubules are polymerized within the ciliary cap, axoneme maturation (the incorporation of axonemal dyneins and other axonemal proteins) may occur within the cytoplasmic compartment, as has been proposed (Avidor-Reiss and Leroux, 2015).

**Figure 6:**
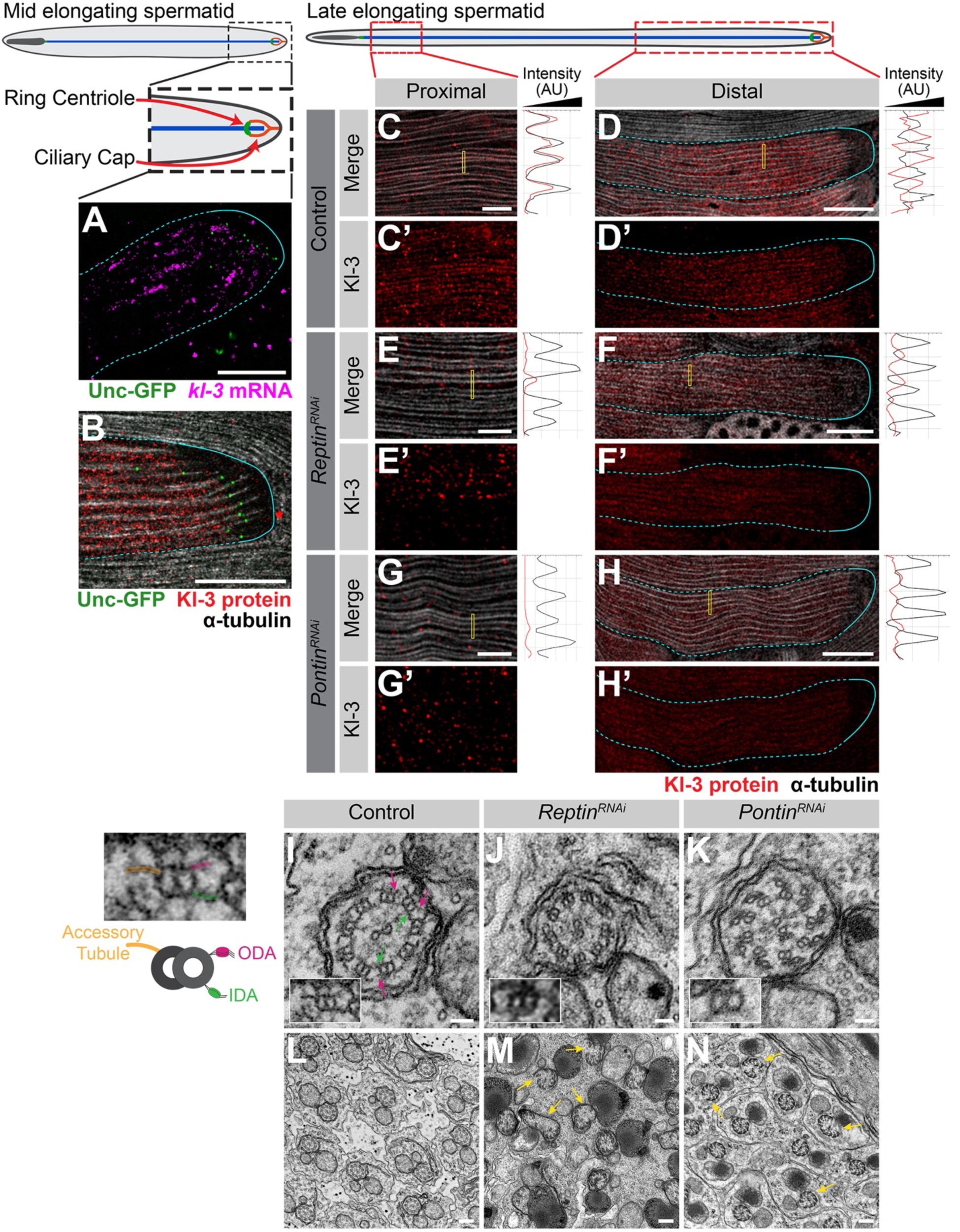
kl-granule formation and localization are required for cytoplasmic cilia maturation. **(A)** smFISH against *kl-3* in flies expressing Unc-GFP. *kl-3* (magenta), Unc-GFP (ring centriole, green), transition zone (yellow arrows), and spermatid cyst (cyan, dashed line: cytoplasmic region, solid line: compartmentalized region). Bar: 20μm. **(B)** Kl-3-3X FLAG protein in flies expressing Unc-GFP. Kl-3 (red), Unc-GFP (ring centriole, green), α-tubulin (white), transition zone (yellow arrows), and spermatid cyst (cyan, dashed line: cytoplasmic region, solid line: compartmentalized region). Bar: 20μm. **(C – H)** Kl-3-3X FLAG protein expression in control (C and D), *rept* RNAi (*bam-gal4>UAS-rept*^*KK105732*^, E and F) and *pont* RNAi (*bam-gal4>UAS-pont*^*KK101103*^, G and H) proximal (C, E and G) and distal (D, F and H) regions of late elongating spermatids. Kl-3 (red), α-tubulin-GFP (white), transition zone (yellow arrows), and spermatid cyst (cyan, dashed line: cytoplasmic region, solid line: compartmentalized region). Intensity plots are shown for the regions within the yellow rectangles. Bar: 5μm (C, E and G) or 25μm (D, F and H). **(I – N)** TEM images of control (I and L), *rept* RNAi (*bam-gal4>UAS-rept*^*KK105732*^, J and M) and *pont* RNAi (*bam-gal4>UAS-pont*^*KK101103*^, K and N) axonemes. Pink arrows: ODA, green arrows: IDA, yellow arrows: broken axonemes broken axonemes. The control single doublet enlarged image is duplicated to the left of the figure and colored to match the diagram. Bar: 50nm (I – K) or 200nm (L – N).

Detailed examination of Kl-3 protein within the elongating spermatid cysts provided insights into where Kl-3 protein may be translated and incorporated into the growing axoneme (Fig. 6 C and D). At the distal end of the cyst, where the kl-granules are concentrated, Kl-3 protein was predominantly observed in the cytoplasm, while being excluded from the axonemal microtubules (Fig. 6D, see the right panel for intensity plot showing mutually exclusive localization of microtubules and Kl-3). This suggests that Kl-3 protein at the distal end may represent the pool of newly translated Kl-3 before it is incorporated into the axoneme, which is also consistent with the presence of kl-granules at this location. In contrast to the distal end, Kl-3 protein was observed to colocalize with axonemal microtubules at the proximal end (Fig. 6D, see the right panel for intensity plot showing colocalization of microtubules and Kl-3), suggesting that Kl-3 protein has been successfully incorporated into the axoneme. These results suggest that Kl-3 protein is translated at the distal end, where the kl-granules localize, and that the diffuse cytoplasmic Kl-3 protein is the newly synthesized pool, which is subsequently incorporated into the axoneme.

Following RNAi mediated knockdown of *rept* or *pont*, which prevents kl-granule formation (Fig. 4) and drastically reduces Kl-3 protein levels (Fig. 5 K), we still observed Kl-3 protein in the cytoplasm at the distal end (Fig. 6 F and H), although at a much reduced level. However, Kl-3 protein was never observed to colocalize with the axonemal microtubules at the proximal end upon *rept* or *pont* RNAi (Fig. 6 E and G), suggesting that Rept and Pont are required for incorporation of Kl-3 into the axoneme.

Consistent with this notion, transmission electron microscopy (TEM) revealed that the ODAs and IDAs are largely absent from the axonemes following *rept* or *pont* RNAi. (Fig. 6 I – K). Additional gross axonemal defects (e.g. broken axonemes) were present in the RNAi conditions (Fig. 6 L – N), suggesting additional impairments to axoneme assembly. These results suggest that localized mRNA translation via formation of the kl-granules is required for axonemal dynein motor proteins to incorporate into the axoneme.

## Discussion

Cytoplasmic cilia have been found in organisms as diverse as *Plasmodium* and humans (Avidor-Reiss et al., 2017; Avidor-Reiss and Leroux, 2015; Dawson and House, 2010; Fawcett et al., 1970; Sinden et al., 1976; Tates, 1971; Tokuyasu, 1975). While it has been proposed that axoneme maturation proceeds through the direct incorporation of axonemal proteins from the cytoplasm, this model remained untested (Avidor-Reiss and Leroux, 2015). Our study provides the first insights into the mechanism of cytoplasmic cilia formation. Our results show that localized translation of axonemal dynein mRNAs facilitates the maturation of cytoplasmic cilia by allowing for the efficient incorporation of axonemal dynein proteins into bare axonemal microtubules directly from the cytoplasm.

### Mechanism for cytoplasmic cilia maturation

It has been proposed that cytoplasmic cilia assemble in two steps (Avidor-Reiss and Leroux, 2015): first, microtubules are polymerized within a small compartmentalized region of the cilia, then, as the bare microtubules are displaced from this region, axonemal proteins are incorporated directly from the cytoplasm during the maturation step. Previous studies that have shown that IFT, the process used by traditional compartmentalized cilia to ferry axonemal proteins into the ciliary compartment, is dispensable for *Drosophila* spermiogenesis, and that the genomes of some other organisms known to form cytoplasmic cilia (e.g. *Plasmodium*) do not encode IFT and/or transition zone proteins (Avidor-Reiss and Leroux, 2015; Breslow et al., 2013; Briggs et al., 2004; Han et al., 2003; Hoeng et al., 2008; Kee et al., 2012; Lin et al., 2013; Sarpal et al., 2003). These studies led to the notion that maturation of cytoplasmic cilia ought to happen in the cytoplasm, although direct evidence has been lacking.

Our study, which identified a novel RNP granule, the kl-granule, composed of axonemal dynein heavy chain mRNAs and the proteins Rept and Pont, provides the first molecular insights into cytoplasmic cilia maturation. Our results show that axonemal dynein heavy chain mRNAs (*kl-3, kl-5, kl-2*, and *Dhc98D*) congress into kl-granules in SCs. We further show that Rept and Pont are required for kl-granule assembly and the proper translation of axonemal dynein mRNAs. We demonstrate that the polarized localization of kl-granule mRNAs within the cytoplasmic compartment promotes localized translation and allows for the incorporation of their encoded proteins into the axoneme, facilitating the maturation step in cytoplasmic cilia assembly (Fig. 7). Our results refine the proposed two step model for cytoplasmic cilia assembly by demonstrating that concentrating axonemal proteins within distal regions of the cytoplasm is critical for maturation. Thus, axoneme maturation proceeds in a stepwise fashion, allowing for the efficient assembly of this very long cilia. This model implies that the proximal region of the axoneme should be more mature than the distal region, a notion that is supported by previous studies that looked at axoneme ultrastructure and tubulin dynamics within the axoneme (Noguchi et al., 2011; Sinden et al., 2010; Tokuyasu, 1975).

**Figure 7:**
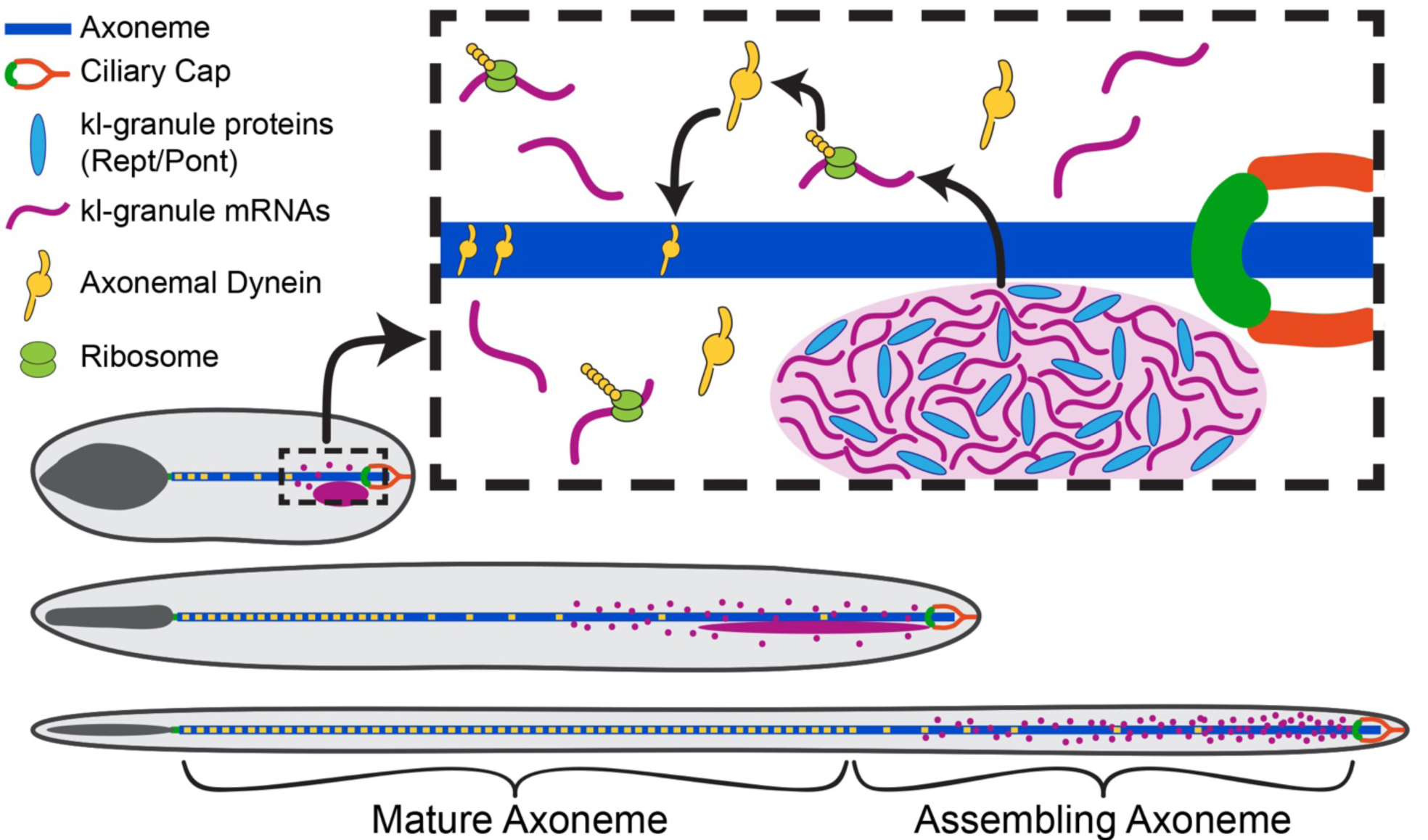
Model for cytoplasmic cilia maturation. The kl-granule (light purple) localizes immediately proximal to the ciliary cap (orange) and ring centriole (green) within the cytoplasmic compartment. Constituent mRNAs (purple) are locally translated (ribosomes, lime green) and their proteins (axonemal dyneins, yellow) are incorporated into the axoneme (blue) as the microtubules are displaced from the ciliary cap. In this way, cytoplasmic cilia maturation is progressive with axonemal proteins being added to the bare microtubules as elongation proceeds.

### Function of Reptin and Pontin in dynein assembly

A wide range of functions have been assigned to Rept- and Pont-containing complexes including roles in chromatin remodeling, transcription regulation, DNA repair and ribosome assembly (Mao and Houry, 2017). They can act alone, together or as part of larger complexes (Huen et al., 2010; Kakihara and Saeki, 2014). Among these, previous studies have proposed that Rept and Pont are dynein arm preassembly factors – chaperones that take individual dynein motor subunits (i.e. the heavy, intermediate, and light chain proteins) and stabilize and assemble them into a motor unit in the cytoplasm that is then ferried into the cilia for incorporation (Desai et al., 2018; Fabczak and Osinka, 2019; Fowkes and Mitchell, 1998). These assembly factors include R2TP and R2TP-like complexes (which include Rept (RUVBL2) and Pont (RUVBL1)) in association with dynein axonemal assembly factors (DNAAFs) (Fabczak and Osinka, 2019). While previous studies have clearly demonstrated that Rept, Pont, R2TP, and DNAAFs are needed for axonemal dynein protein stability and incorporation (Huizar et al., 2018; Li et al., 2017; Liu et al., 2019; Yamaguchi et al., 2018; Zhao et al., 2013; Zur Lage et al., 2018), our study is the first to demonstrate involvement of axonemal dynein mRNAs with these complexes, showing that Rept and Pont are required for axonemal dynein mRNAs to localize to the kl-granules. It remains unknown whether the Rept- and Pont-associated dynein arm preassembly complexes reported in previous studies also contain dynein mRNAs. However, important differences exist between these preassembly complexes and the kl-granules. Firstly, while *kl-3* mRNA is present in the kl-granules, no puncta are observed for Kl-3 protein, indicating that Kl-3 protein does not concentrate within the kl-granules (or another granule) as dyneins do in the dynein preassembly complexes reported in other systems (Dafinger et al., 2018; Huizar et al., 2018). Additionally, dynein preassembly complexes were found to contain proteins (e.g. Wdr78 (Huizar et al., 2018)), where the *Drosophila* homolog (*Dic61B*) mRNAs are not constituents of the kl-granules (Fig. S4, see below). Therefore, the kl-granule may be a novel adaptation of a Rept and Pont containing dynein arm assembly complex specifically found in cytoplasmic cilia and is distinct from its role as a dynein preassembly factor in other systems.

It is likely that additional protein components of the kl-granule remain to be discovered. Structural analyses in previous studies have identified mechanisms by which other proteins interact with Rept and Pont (Rivera-Calzada et al., 2017), however, Rept and Pont do not have any RNA binding domains (Mao and Houry, 2017). Therefore, it is likely that additional proteins, not Rept and Pont themselves, physically interact with constituent mRNAs for kl-granule formation. Our data also supports the existence of additional proteins governing kl-granule dynamics. For example, as spermatids elongate, the ODA and IDA mRNAs separate slightly from each other while remaining polarized at the distal end (Fig. 2). Moreover, in the absence of Rept and Pont, the IDA mRNAs are still able to congress at the distal end of the elongating spermatid cyst, after failing to form kl-granules in SCs. In contrast, localization of the ODA mRNAs entirely depends on Rept and Pont, as ODA mRNAs remain diffuse throughout spermatogenesis following *rept* or *pont* RNAi. Finally, Pont more strongly colocalizes with the ODA mRNAs within the kl-granule (Fig. 3 F), which altogether suggests that there are additional proteins that can sort and specify the fate of these kl-granule mRNAs both alongside or in the absence of Rept and Pont. The identity of these additional proteins is the subject of further study. In addition, determining the involvement of the other dynein arm preassembly factors is of particular interest, especially considering the existence of multiple dynein arm assembly complexes that have been shown to differentially regulate IDA and ODA assembly (Fabczak and Osinka, 2019; Yamaguchi et al., 2018). It is also appealing to posit the existence of testis-specific factors which may help to distinguish the role of Rept and Pont in cytoplasmic cilia formation from its role in the assembly of other cellular bodies.

### Purpose of mRNA localization to kl-granules

Interestingly, we found that not all mRNAs for axonemal/spermiogenesis proteins localize to the kl-granules (Fig. S4). mRNAs for other axonemal proteins (the dynein intermediate chain *Dic61B*, the dynein heavy chain *CG3339*, and the ODA docking complex component *CG17083* (Zur Lage et al., 2019)) as well as mRNAs for other Y-linked transcripts (*CCY* and *PPR-Y* (Carvalho et al., 2001)) and a non-axonemal spermatid protein (*fzo* (Hales and Fuller, 1997)) did not localize to the kl-granules. Instead they remain evenly distributed throughout the SC cytoplasm, despite also being important for sperm maturation (Fig. S4). Additionally, we previously reported that mRNAs for the Y-liked gene *ORY* also gather in cytoplasmic RNA granules in late SCs (Fingerhut et al., 2019), however, these RNA granules are distinct from the kl-granule (Fig. S4 G).

In particular, it is intriguing that mRNA for *Dic61B*, an IDA intermediate chain that needs to bind to the IDA heavy chains Kl-2 and Dhc98D, is located differently (diffusely) within the spermatid cyst. Dynein preassembly is believed to be important for dynein protein stability and a prerequisite for axonemal incorporation (Fabczak and Osinka, 2019; Fowkes and Mitchell, 1998). An intriguing possibility is that temporal/spatial regulation of dynein mRNAs plays a role in helping the ordered assembly of dynein complexes. It will be of future interest to determine when and where during spermiogenesis dynein complexes are formed in the cytoplasmic cilia as well as what factors are necessary for their formation. A comprehensive understanding of kl-granule mRNAs and proteins would allow for further study into this temporal/spatial regulatory mechanism and a more thorough understanding of how the kl-granules function in the maturation of cytoplasmic cilia.

In summary, our study provides the first insights into the mechanism of cytoplasmic cilia maturation: mRNAs for axonemal dynein motor proteins are localized at the distal end of the axoneme within the cytoplasmic compartment, which allows for efficient maturation of cytoplasmic cilia through localized translation.

## Materials and Methods

### Fly husbandry

All fly stocks were raised on standard Bloomington medium at 25°C, and young flies (1- to 5-day-old adults) were used for all experiments. Flies used for wildtype experiments were the standard lab wildtype strain *yw* (*y*^*1*^*w*^*1*^). The following fly stocks were used: *bam-GAL4:VP16* (BDSC:80579), *UAS-kl-3*^*TRiP.HMC03546*^ (BDSC:53317), *UAS-kl-5*^*TRiP.HMC03747*^ (BDSC:55609), *UAS-Dhc98D*^*TRiP.HMC06494*^ (BDSC:77181), and C(1)RM/C(1;Y)6, *y*^*1*^*w*^*1*^*f*^*1*^/0 (BDSC:9460) were obtained from the Bloomington Stock Center (BDSC). *UAS-kl-2*^*GC8807*^ (VDRC:v19181), *UAS-rept*^*KK105732*^ (VDRC:v103483), and *UAS-pont*^*KK101103*^ (VDRC:v105408) were obtained from the Vienna *Drosophila* Resource Center (VDRC). *unc-GFP* (GFP-tagged *unc* expressed by the endogenous promoter) and Ub-α-tubulin84B-GFP were a gift of Cayentano Gonzalez (Baker et al., 2004; Rebollo et al., 2004) and *bam-gal4* was a gift of Dennis McKearin (Chen and McKearin, 2003). The *kl-3-FLAG* strain was constructed using CRISPR mediated knock-in of a 3X-FLAG tag at the C-terminus of *kl-3* as previously described (Fingerhut et al., 2019).

### Single molecule RNA fluorescent *in situ* hybridization

All solutions used for RNA FISH were RNase free. Testes from 2-3 day old flies were dissected in 1X PBS and fixed in 4% formaldehyde in 1X PBS for 30 minutes. Then testes were washed briefly in 1X PBS and permeabilized in 70% ethanol overnight at 4°C. Testes were briefly rinsed with wash buffer (2X saline-sodium citrate (SSC), 10% formamide) and then hybridized overnight at 37°C in hybridization buffer (2X SSC, 10% dextran sulfate (Sigma, D8906), 1mg/mL E. coli tRNA (Sigma, R8759), 2mM Vanadyl Ribonucleoside complex (NEB S142), 0.5% bovine serum albumin (BSA, Ambion, AM2618), 10% formamide). Following hybridization, samples were washed three times in wash buffer for 20 minutes each at 37°C and mounted in VECTASHIELD with DAPI (Vector Labs). Images were acquired using an upright Leica TCS SP8 confocal microscope with a 63X oil immersion objective lens (NA = 1.4) and processed using Adobe Photoshop and ImageJ software. Fluorescently labeled probes were added to the hybridization buffer to a final concentration of 100nM. Probes against *kl-3, kl-5, kl-2, Dhc98D, CG3339, Dic61B, CG17083, CCY, PPR-Y, ORY* and *fzo* mRNAs were designed using the Stellaris® RNA FISH Probe Designer (Biosearch Technologies, Inc.) available online at www.biosearchtech.com/stellarisdesigner. Each set of custom Stellaris® RNA FISH probes was labeled with Quasar 670, Quasar 570 or Fluorescein (Table S1).

For strains expressing GFP (e.g. unc-GFP or Ub-α-tubulin84B-GFP), the overnight permeabilization in 70% ethanol was omitted.

### Immunofluorescence staining

Testes were dissected in 1X PBS, transferred to 4% formaldehyde in 1X PBS and fixed for 30 minutes. Testes were then washed in 1X PBST (PBS containing 0.1% Triton-X) for at least 60 minutes followed by incubation with primary antibodies diluted in 1X PBST with 3% BSA at 4°C overnight. Samples were washed for at least 1 hour in 1X PBST, incubated with secondary antibody in 1X PBST with 3% BSA at 4°C overnight, washed as above, and mounted in VECTASHIELD with DAPI (Vector Labs). Images were acquired using an upright Leica TCS SP8 confocal microscope with a 63X oil immersion objective lens (NA = 1.4) and processed using Adobe Photoshop and ImageJ software.

The following primary antibodies were used: anti–α-tubulin (1:100; mouse, Sigma-Aldrich T6199), anti-FLAG (1:500; rabbit, Invitrogen PA1-984B), anti-Reptin (1:200; rabbit, gift of Andrew Saurin (Diop et al., 2008)), anti-Pontin (1:200; guinea pig, this study), Phalloidin-Alexa546 or 488 (1:200; ThermoFisher A22283 or A12379). The Pontin antibody was generated by injecting a peptide (CKVNGRNQISKDDIEDVH, targeting 18aa from the c terminal end of Pontin) in guinea pigs (Covance). Alexa Fluor-conjugated secondary antibodies (Life Technologies) were used at a dilution of 1:200.

A modified version of Stefanini’s fixative (4% formaldehyde, 0.18% w/v Picric Acid (Ricca Chemical 5860), 0.3M PIPES pH7.5 (Alfa Aesar J63617), 0.05% Tween-20) was used in order to detect Kl-3 (Muller, 2008). No signal was detectable using traditional formaldehyde fixation.

### Immunofluorescence staining with single molecule RNA fluorescent *in situ* hybridization

To combine immunofluorescent staining with smFISH, testes from 2-3 day old flies were dissected in 1X PBS and fixed in 4% formaldehyde in 1X PBS for 30 minutes. Then testes were washed briefly in PBS and permeabilized in 70% ethanol overnight at 4°C (unless from a strain expressing GFP, in which case this step was omitted). Testes were then washed with 1X PBS and blocked for 30 minutes at 37°C in blocking buffer (1X PBS, 0.05% BSA, 50μg/mL E. coli tRNA, 10mM Vanadyl Ribonucleoside complex, 0.2% Tween-20). Primary antibodies were diluted in blocking buffer and incubated at 4°C overnight. The testes were washed with 1X PBS containing 0.2% Tween-20, re-blocked for 5 minutes at 37°C in blocking buffer and incubated 4°C overnight in blocking buffer containing secondary antibodies. Then testes were washed with 1X PBS containing 0.2% Tween-20 and re-fixed for 10 minutes before continuing the smFISH starting from the brief rinse with wash buffer.

### RT-qPCR

Total RNA from testes (50 pairs/sample) was extracted using TRIzol (Invitrogen) according to the manufacturer’s instructions. 1μg of total RNA was reverse transcribed using SuperScript III® Reverse Transcriptase (Invitrogen) followed by qPCR using *Power* SYBR Green reagent (Applied Biosystems) on a QuantStudio 6 Real-Time PCR system (Applied Biosystems). Primers for qPCR were designed to amplify only mRNA. The genes analyzed by qPCR are all predicted to contain megabase sized introns, and primers were designed to span these large introns such that a product would be detect only if the intron had been spliced out (Fingerhut et al., 2019). Relative expression levels were normalized to GAPDH and cross-sibling controls. All reactions were done in technical triplicates with at least two biological replicates. Graphical representation was inclusive of all replicates. Primers used are listed in Table S1.

### Western blot

Testes (40 pairs/sample) were dissected in Schneider’s media at room temperature within 30 minutes, the media was removed and the samples were frozen at −80°C until use. After thawing, testes were then lysed in 200uL of 2X Laemmli Sample Buffer + βME (BioRad 161-0737). For Kl-3, samples were separated on a NuPAGE Tris-Acetate gel (3-8%, 1.5mm, Invitrogen) and for Rept and Pont, samples were separated on a Novex Tris-Glycine gel (10%, 1mm, Invitrogen) with the appropriate running buffer in a Xcell SureLock mini-cell electrophoresis system (Invitrogen). For Kl-3, proteins were transferred using the XCell II blot module (Invitrogen) onto polyvinylidene fluoride (PVDF) membrane (Immobilon-P, Millipore) using NuPAGE transfer buffer (Invitrogen) without added methanol. For Rept and Pont, transfer buffer contained 20% methanol. Membranes were blocked in 1X TBST (0.1% Tween-20) containing 5% nonfat milk, followed by incubation with primary antibodies diluted in 1X TBST containing 5% nonfat milk. Membranes were washed with 1X TBST, followed by incubation with secondary antibodies diluted in 1X TBST containing 5% nonfat milk. After washing with 1X TBST, detection was performed using the Pierce® ECL Western Blotting Substrate enhanced chemiluminescence system (Thermo Scientific). Primary antibodies used were anti–α-tubulin (1:2,000; mouse, Sigma-Aldrich T6199), anti-FLAG (1:2,500; mouse, Sigma-Aldrich F1804), anti-Reptin (1:2000; rabbit, gift of Andrew Saurin), anti-Pontin (1:2000; guinea pig, this study), anti-Vasa (1:3000; rabbit, Santa Cruz Biotechnology D-260). The secondary antibodies were horseradish peroxidase (HRP) conjugated goat anti-mouse IgG, anti-rabbit IgG, or anti-guinea pig IgG (1:10,000; Abcam).

### Phase contrast microscopy

Seminal vesicles were dissected in 1X PBS and transferred to slides for live observation by phase contrast on a Leica DM5000B microscope with a 40X objective (NA = 0.75) and imaged with a QImaging Retiga 2000R Fast 1394 Mono Cooled camera. Images were adjusted in Adobe Photoshop.

### Transmission electron microscopy

Testes were fixed for one hour or overnight (at 4°C) with 2.5% glutaraldehyde in 0.1M Sorensen’s buffer, pH7.4. Samples were rinsed twice for 5 minutes each with 0.1 M Sorensen’s buffer and post fixed for one hour in 1 % osmium tetroxide in 0.1 M Sorensen’s buffer. Next, testes were rinsed twice in double distilled water for 5 minutes each and *en bloc* stained with 2 % uranyl acetate in double distilled water for one hour. The samples were them dehydrated in increasing concentrations of ethanol, rinsed with acetone, and embedded in Epon epoxy resin. Thin sections were mounted on Formvar/carbon-coated slotted grids and post-stained with uranyl acetate and lead citrate. Samples were examined on a JEOL1400 transmission electron microscope and images captured using a sCMOS XR401 custom engineered optic camera by AMT (Advanced Microscopy Techniques Corp.).

### Online supplemental material

Fig. S1 shows efficiency of RNAi knockdown of *kl-3, kl-5, kl-2*, and *Dhc98D* by smFISH and lack of dependence upon a single one of those transcripts for kl-granule formation (related to Fig. 1). Fig. S2 shows that RNAi of *rept* or *pont* results in efficient knockdown of both (related to Fig. 4). Fig. S3 shows the sterility phenotype of *kl-3, kl-5, kl-2*, and *Dhc98D* RNAi flies (related to Fig. 5). Fig. S4 shows smFISH for other axonemal, Y-linked, and spermatid-essential transcripts (related to Discussion).

## Abbreviations

SC: spermatocyte
IFT: intraflagellar transport
ODA: Outer dynein arm
IDA: Inner dynein arm
RNP: Ribonucleoprotein
smFISH: single molecule RNA fluorescent in situ hybridization
IF: Immunofluorescent
IC: Individualization complex
TEM: Transmission light microscopy

## Acknowledgements

We thank Drs. Dennis McKearin, Cayentano Gonzalez, and Andrew Saurin, the Bloomington Stock Center, and the Vienna *Drosophila* Resource Center for reagents. We thank Sasha Meshinchi and the University of Michigan Microscopy Core for help with EM experiments. We thank the Yamashita laboratory and Drs. Sue Hammoud and Joshua Bembenek for discussion and comments on the manuscript, Drs. Tomer Avidor-Reiss and Tony Mahowald for helpful suggestions, and Dr. Jiandie Lin for sharing equipment. This work was supported by the Howard Hughes Medical Institute (to YMY) and the NIH Cellular and Molecular Biology Training Grant T32-GM007315 (to JMF). The authors declare no competing financial interests.

## Author Contributions

J.M. Fingerhut conceived the project and conducted experiments. J.M. Fingerhut and Y.M. Yamashita designed experiments, analyzed the data, and wrote the manuscript.

## Figures

**Figure S1:**
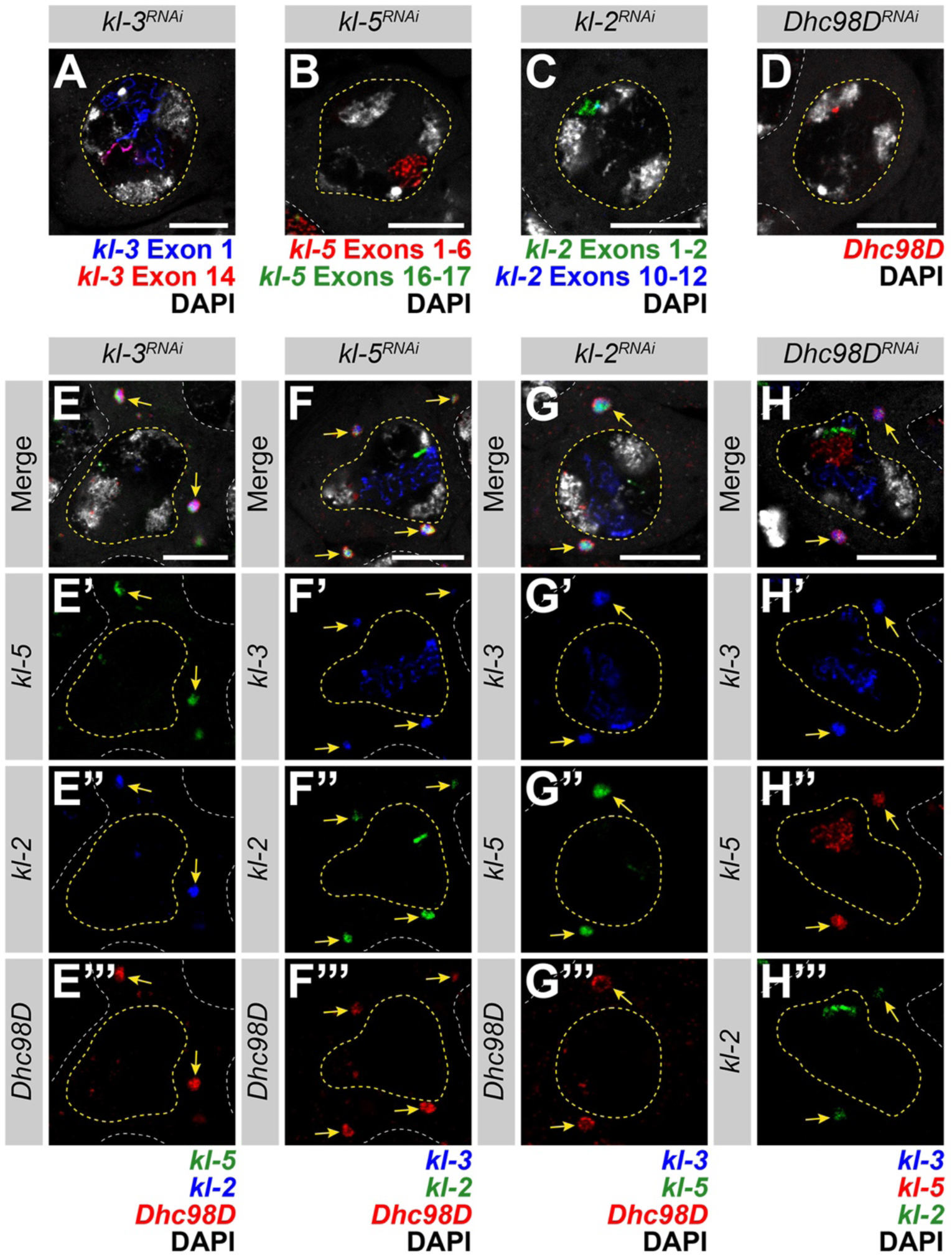
kl-granule formation is not dependent upon any one mRNA constituent. **(A – D)** smFISH against each known kl-granule mRNA constituent following RNAi of that constituent shows successful knockdown (no remaining cytoplasmic signal). Note that we use multiple smFISH probe sets for some mRNAs targeted against different regions of the transcript (see Table S1). (A) *kl-3* exon 1 (blue), *kl-3* exon 14 (red) and DAPI (white). (B) *kl-5* exons 1-6 (red), *kl-5* exons 16-17 (green) and DAPI (white). (C) *kl-2* exons 1-2 (green), *kl-2* exons 10-12 (blue) and DAPI (white). (D) *Dhc98D* (red) and DAPI (white). For all, SC nuclei (yellow dashed line) and neighboring SC nuclei (white dashed line). Bar: 10μm. **(E – H)** smFISH against the other three constituent mRNAs after RNAi of the fourth mRNA. Note that the color used to represent each smFISH probe corresponds to the probe sets in A – D. (E) *kl-5* (green), *kl-2* (blue), *Dhc98D* (red) and DAPI (white). (F) *kl-3* (blue), *kl-5* (green), *Dhc98D* (red) and DAPI (white). (G) *kl-3* (blue), *kl-5* (green), *Dhc98D* (red) and DAPI (white). (H) *kl-3* (blue), *kl-5* (red), *kl-2* (green) and DAPI (white). For all, SC nuclei (yellow dashed line), neighboring SC nuclei (white dashed line), and kl-granules (yellow arrows). Bar: 10μm.

**Figure S2:**
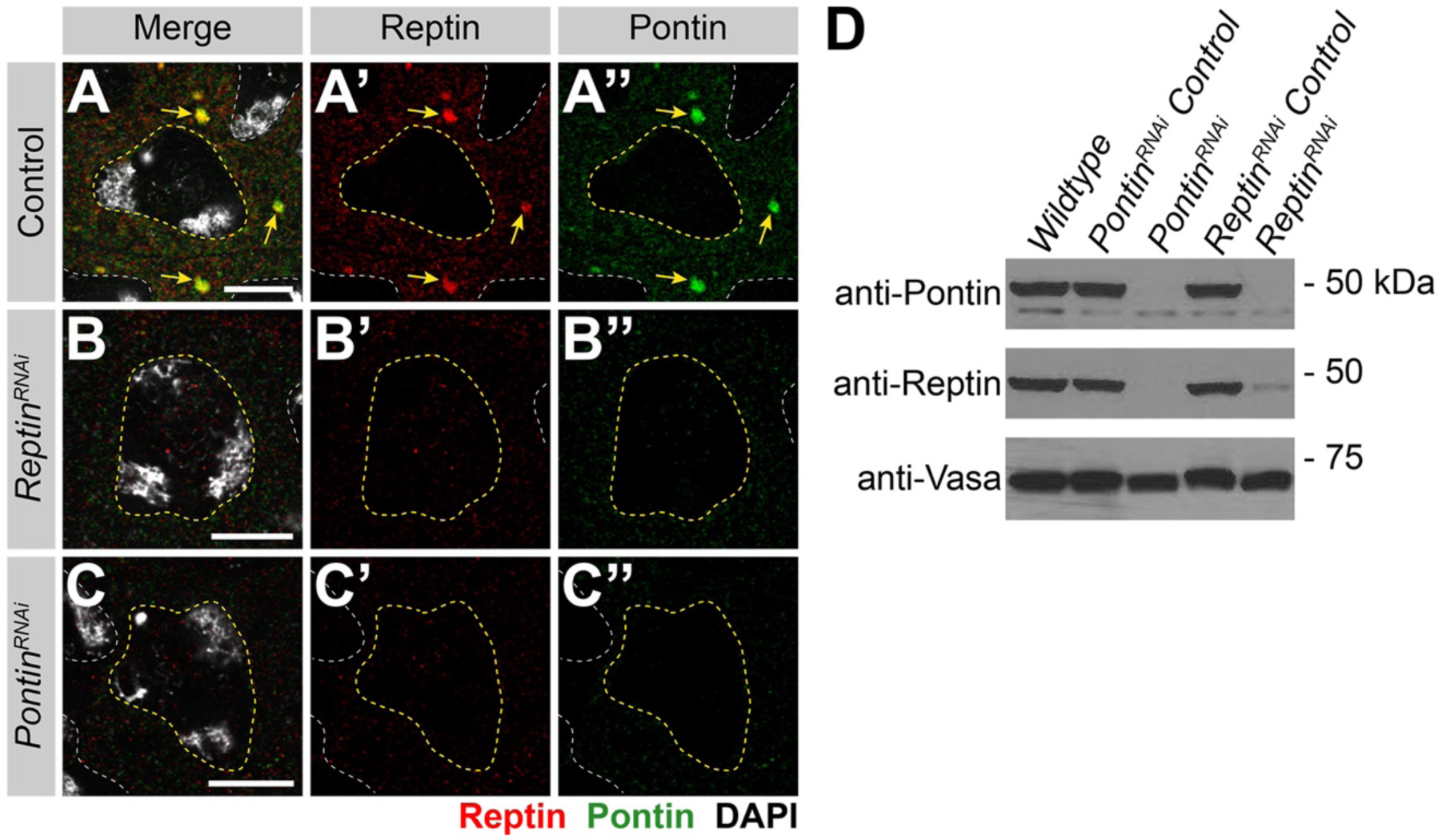
RNAi of *rept* or *pont* results in loss of both proteins. **(A – C)** Rept and Pont protein expression in SCs in control (A), *rept* RNAi (*bam-gal4>UAS-rept*^*KK105732*^, B) or *pont* RNAi (*bam-gal4>UAS-pont*^*KK101103*^, C). Rept (red), Pont (green), DAPI (white), SC nuclei (yellow dashed line), neighboring SC nuclei (white dashed line), kl-granules (yellow arrow). Bar: 10μm. **(D)** Western blot for Pont and Rept in the indicated genotypes.

**Figure S3:**
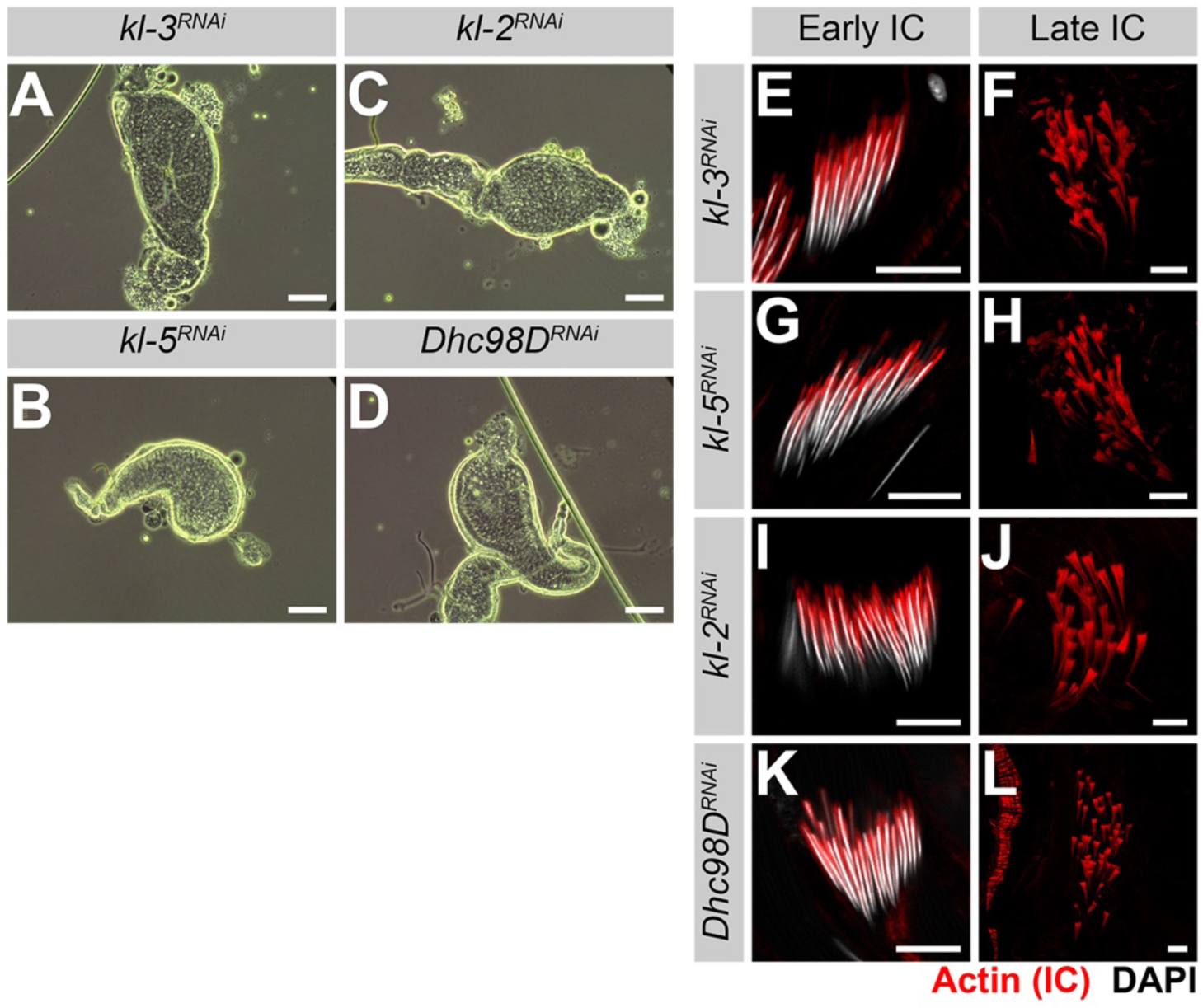
RNAi of *kl-3, kl-5, kl-2* or *Dhc98D* results in the same sterility phenotype seen in *rept* or *pont* RNAi testes. **(A – D)** Phase contrast images of seminal vesicles in *kl-3* RNAi (*bam-gal4>UAS-kl-3*^*TRiP.HMC03546*^, A), *kl-5* RNAi (*bam-gal4>UAS-kl-5*^*TRiP.HMC03747*^, B), *kl-2* RNAi (*bam-gal4>UAS-kl-2*^*GC8807*^, C) and *Dhc98D* RNAi (*bam-gal4>UAS-Dhc98D*^*TRiP.HMC06494*^, D). Bar: 100μm. **(E – L)** Phalloidin staining of early and late ICs in the indicated genotypes. Phalloidin (Actin, red) and DAPI (white). Bar 10μm.

**Figure S4:**
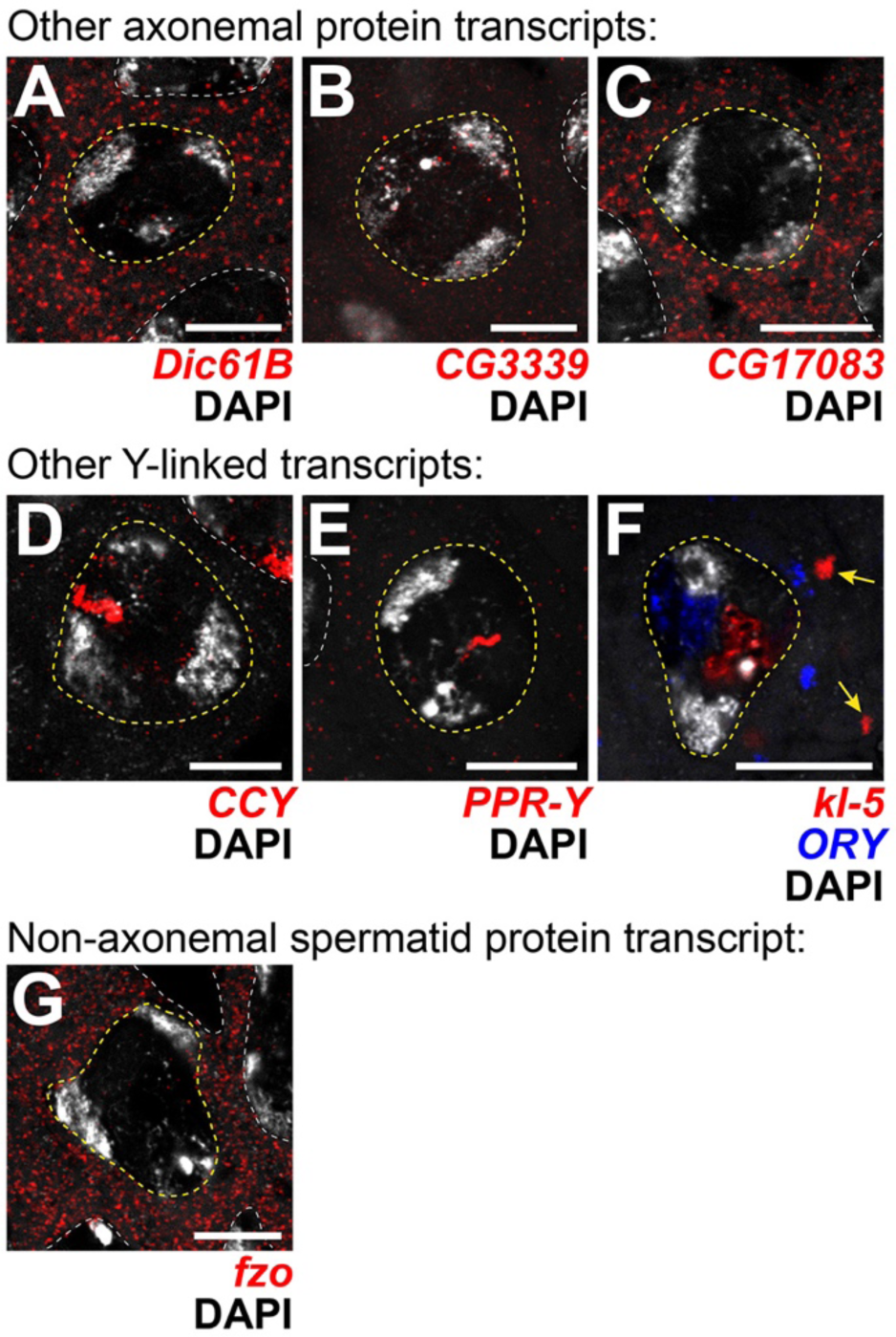
Transcripts for other axonemal, Y-linked, and spermatid proteins don’t localize to kl-granules. **(A – G)** smFISH against other axonemal (A – C), Y-linked (D – F) or spermatid-essential (G) transcripts. (A) *Dic61B* (dynein intermediate chain, red) and DAPI (white). (B) *CG3339* (axonemal dynein heavy chain, red) and DAPI (white). (C) *CG17083* (ODA docking complex, red) and DAPI (white). (D) *CCY* (red) and DAPI (white). (E) *PPR-Y* (red) and DAPI (white). (F) *kl-5* (red), *ORY* (blue) DAPI (white) and kl-granules (yellow arrow). (G) *fzo* (red) and DAPI (white). For all, SC nuclei (yellow dashed line) and neighboring SC nuclei (white dashed line). Bar: 10μm.

**Table S1:**
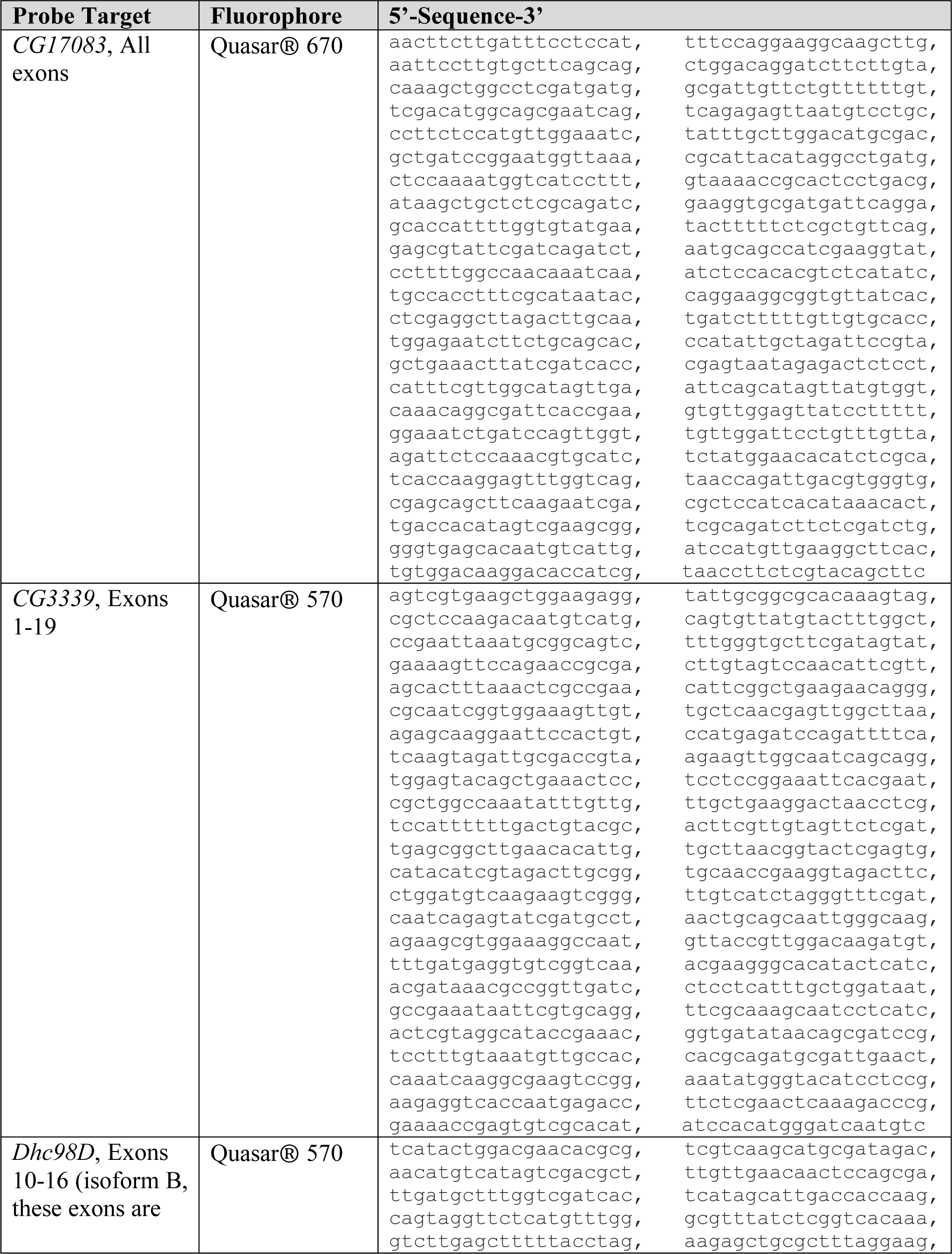
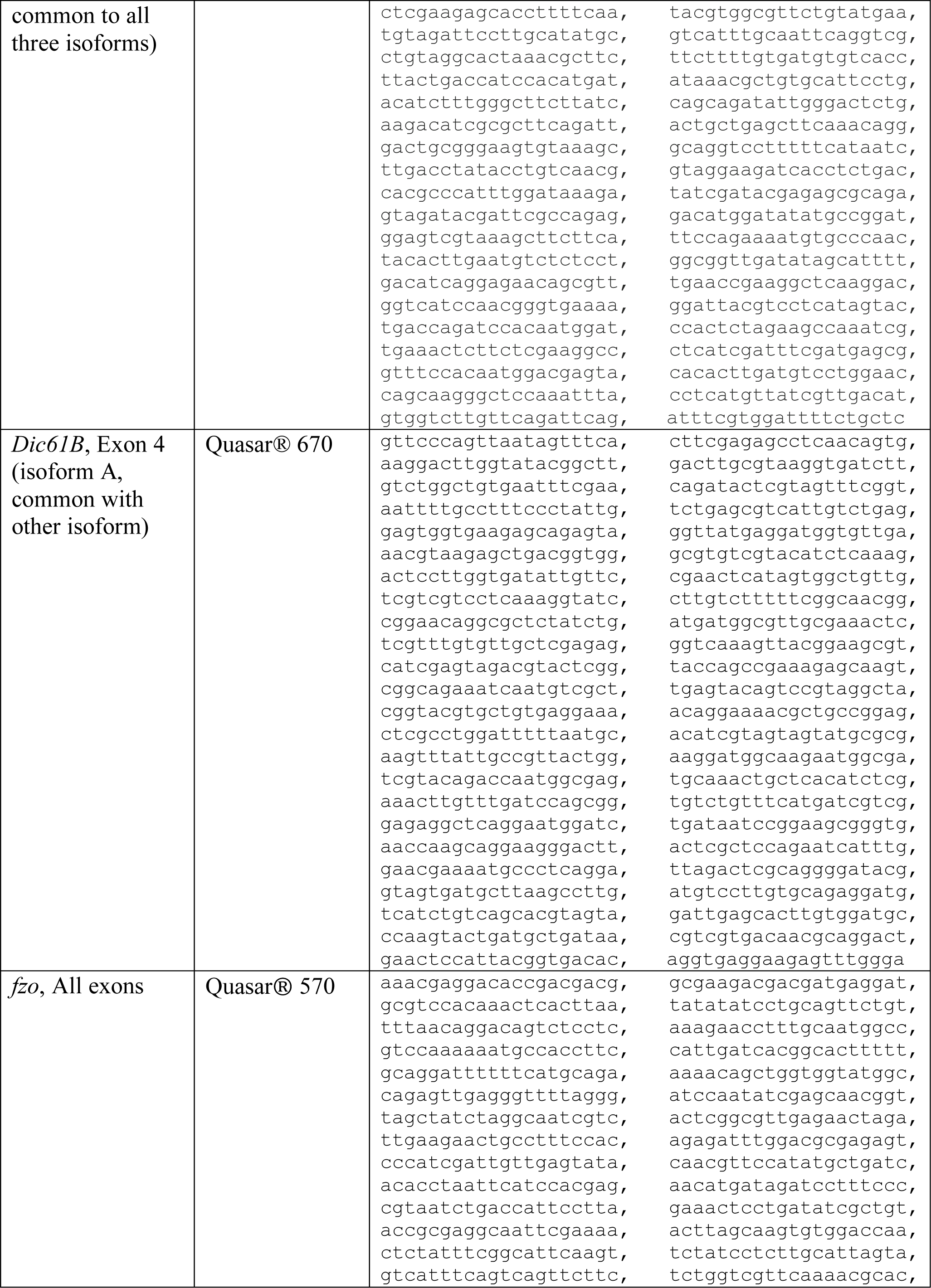
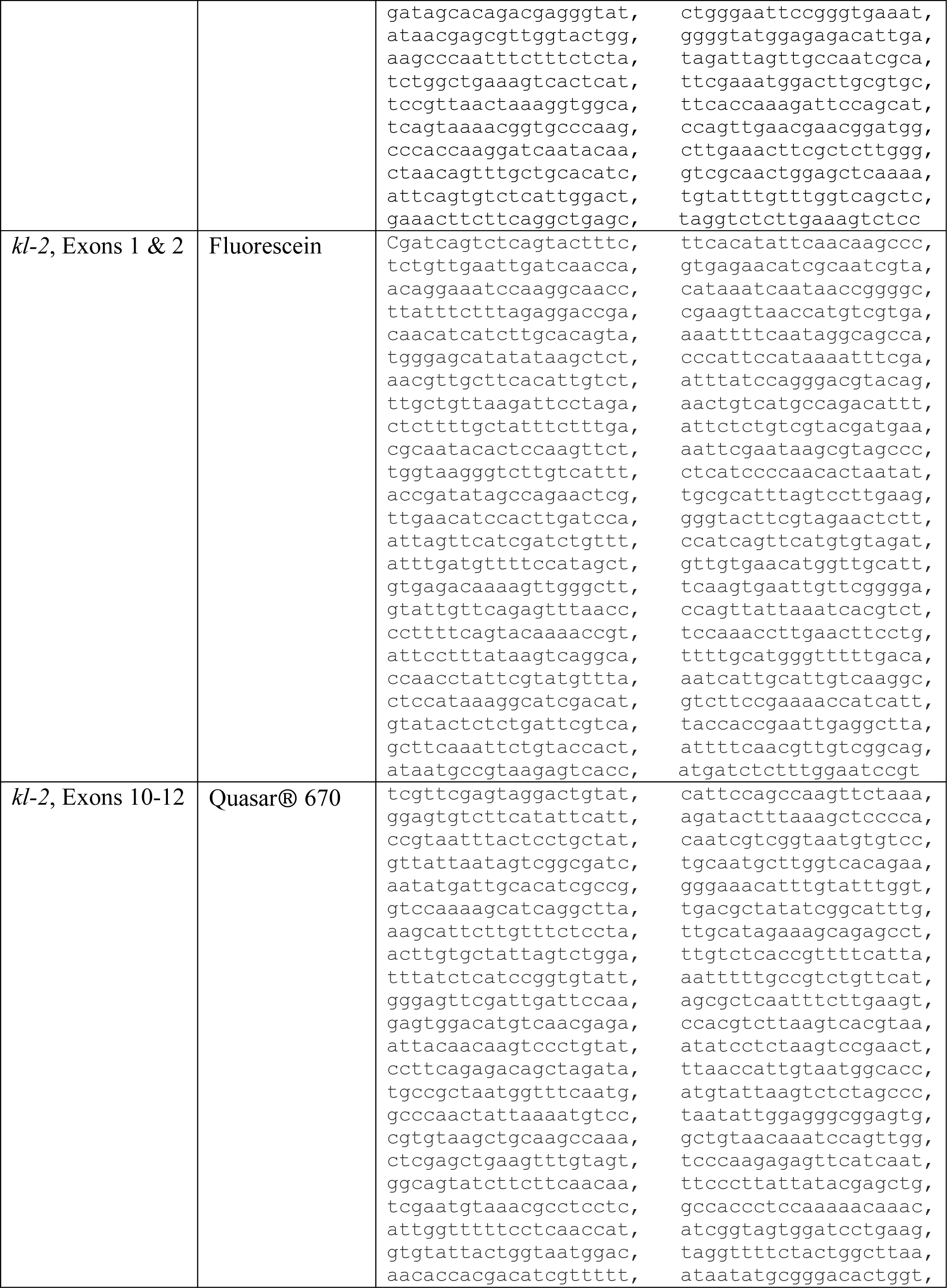
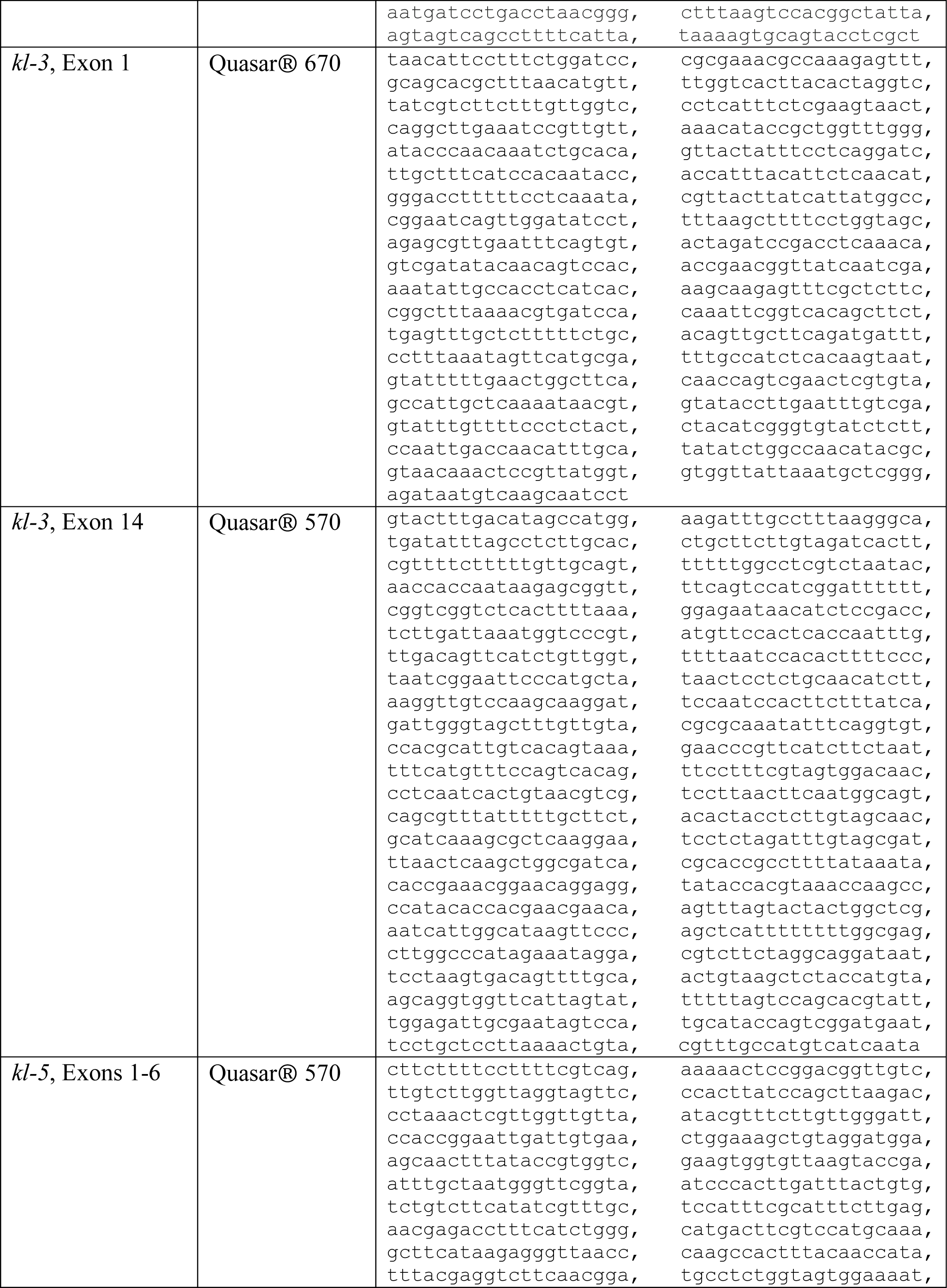
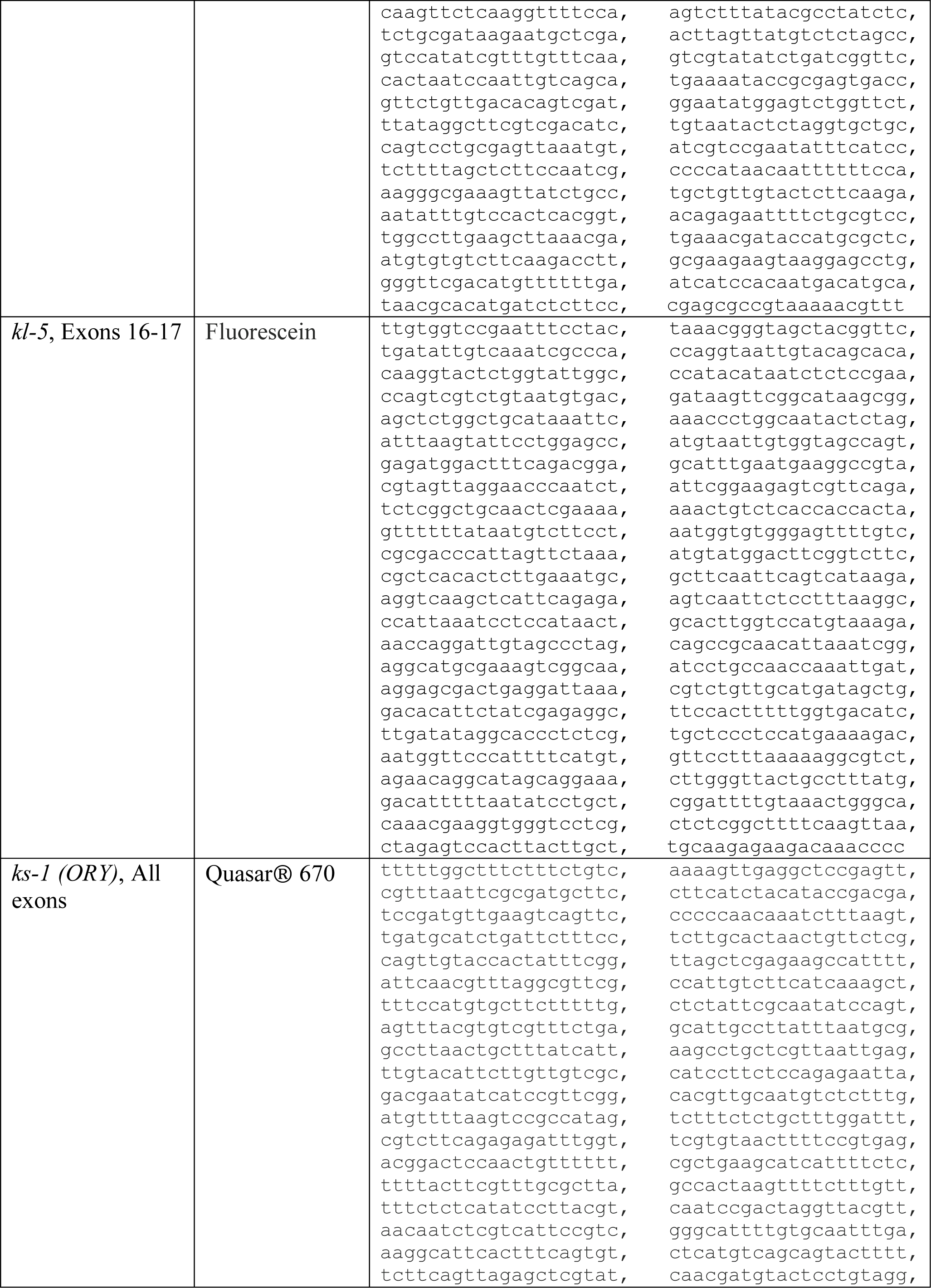
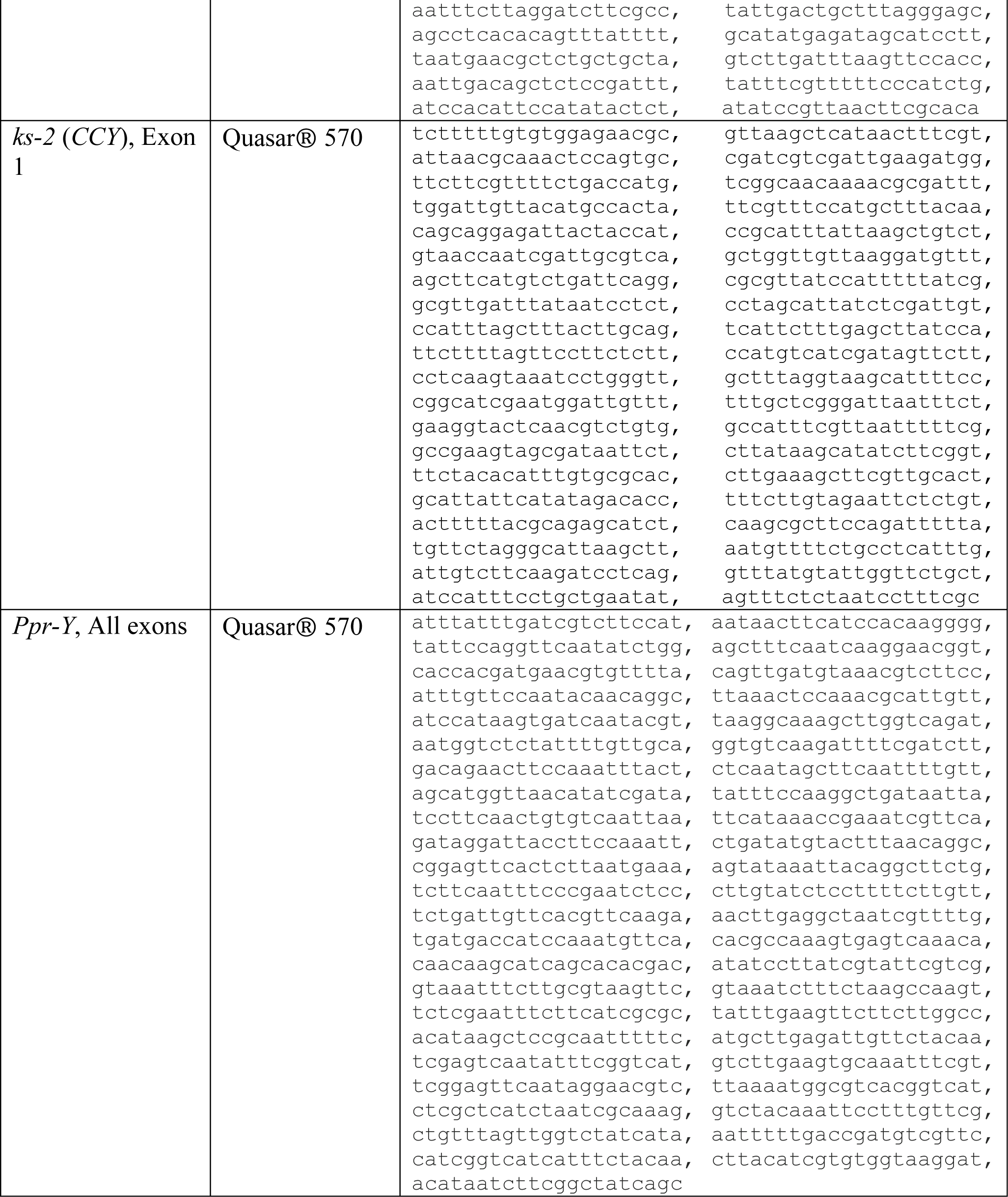
smFISH probes and RT-qPCR primers used in this study.

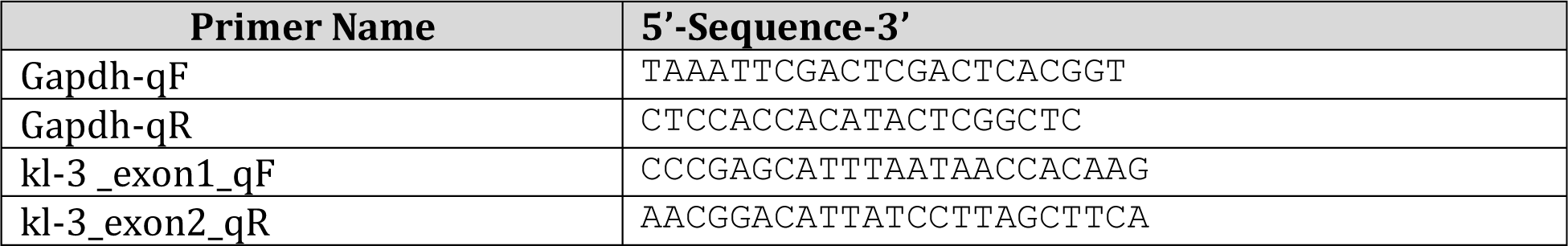
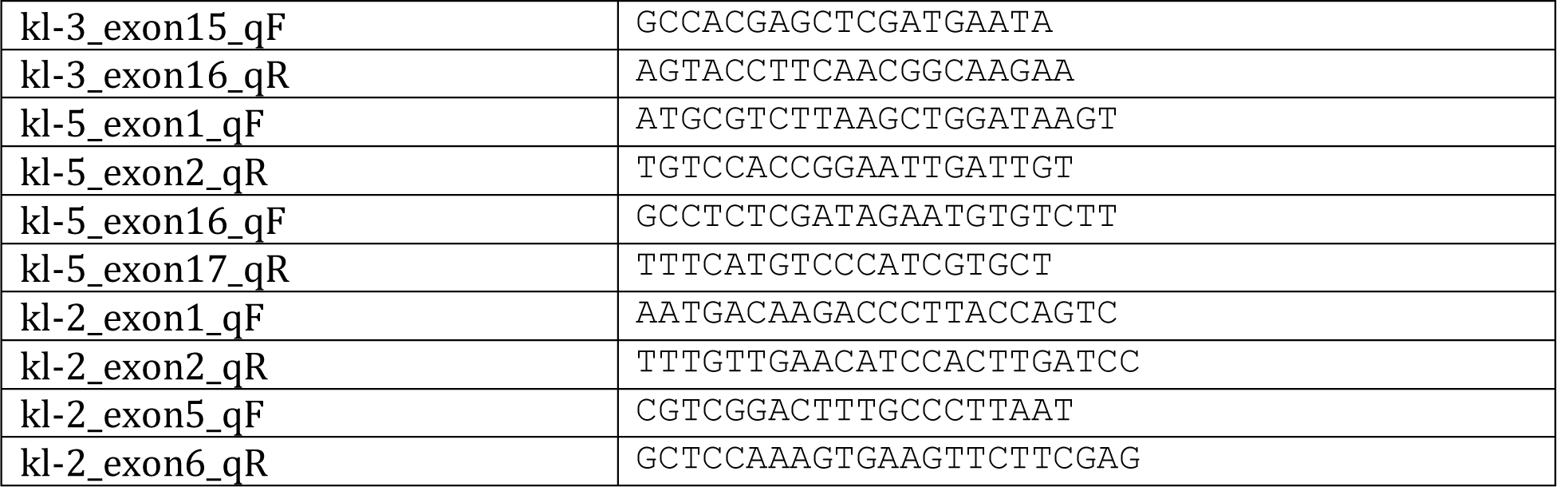

## Notes

### Competing Interest Statement

The authors have declared no competing interest.

